# Transcription shapes 3D chromatin organization by interacting with loop extrusion

**DOI:** 10.1101/2022.01.07.475367

**Authors:** Edward J. Banigan, Wen Tang, Aafke A. van den Berg, Roman R. Stocsits, Gordana Wutz, Hugo B. Brandão, Georg A. Busslinger, Jan-Michael Peters, Leonid A. Mirny

**Author notes:** These authors contributed equally to this work (Edward J. Banigan, Wen Tang, Aafke A. van den Berg). Correspondence (JMP) and (LAM).

## Abstract

Cohesin folds mammalian interphase chromosomes by extruding the chromatin fiber into numerous loops. “Loop extrusion” can be impeded by chromatin-bound factors, such as CTCF, which generates characteristic and functional chromatin organization patterns. It has been proposed that transcription relocalizes or interferes with cohesin, and that active promoters are cohesin loading sites. However, the effects of transcription on cohesin have not been reconciled with observations of active extrusion by cohesin. To determine how transcription modulates extrusion, we studied mouse cells in which we could alter cohesin abundance, dynamics, and localization by genetic ‘knockouts’ of the cohesin regulators CTCF and Wapl. Through Hi-C experiments, we discovered intricate, cohesin-dependent contact patterns near active genes. Chromatin organization around active genes exhibited hallmarks of interactions between transcribing RNA polymerases (RNAPs) and extruding cohesins. These observations could be reproduced by polymer simulations in which RNAPs were “moving barriers” to extrusion that obstructed, slowed, and pushed cohesins. The simulations predicted that preferential loading of cohesin at promoters is inconsistent with our experimental data. Additional ChIP-seq experiments showed that the putative cohesin loader Nipbl is not predominantly enriched at promoters. Therefore, we propose that cohesin is not preferentially loaded at promoters and that the barrier function of RNAP accounts for cohesin accumulation at active promoters. Altogether, we find that RNAP is a new type of extrusion barrier that is not stationary, but rather, translocates and relocalizes cohesin. Loop extrusion and transcription might interact to dynamically generate and maintain gene interactions with regulatory elements and shape functional genomic organization.

**Significance Statement:** Loop extrusion by cohesin is critical to folding the mammalian genome into loops. Extrusion can be halted by CTCF proteins bound at specific genomic loci, which generates chromosomal domains and can regulate gene expression. However, the process of transcription itself can modulate cohesin, thus refolding chromosomes near active genes. Through experiments and simulations, we show that transcribing RNA polymerases (RNAPs) act as “moving barriers” to loop-extruding cohesins. Unlike stationary CTCF barriers, RNAPs actively relocalize cohesins, which generates characteristic patterns of spatial organization around active genes. Our model predicts that the barrier function of RNAP can explain why cohesin accumulates at active promoters and provides a mechanism for clustering active promoters. Through transcription-extrusion interactions, cells might dynamically regulate functional genomic contacts.

## Introduction

The cohesin protein complex organizes mammalian interphase chromosomes by reeling chromatin fibers into dynamic loops in a process known as “loop extrusion” (1–5). While cohesin is bound to chromatin, it can progressively grow chromatin loops until extrusion is obstructed. Obstructions to loop extrusion, such as properly oriented CTCF proteins (6–12), generate characteristic patterns of chromatin organization, such as insulating domains (*e.g*., topologically associating domains, “TADs”) (13–17). Within insulated regions, genomic contacts are enriched, while contacts across CTCF boundaries are suppressed (13–17). Extrusion barriers can thus facilitate or suppress functional interactions, such as enhancer-promoter contacts, which can impact differentiation, disease, and other physiological processes (8, 18–24). Other factors that do not occupy specific genomic positions, such as the replicative helicase MCM, can also act as barriers to loop extrusion (25). These observations raise the question of how chromatin organization by loop-extruding cohesins is shaped by other chromatin-bound factors, some of which may themselves be mobile. We thus investigated how transcription affects loop extrusion and thereby modulates the 3D organization of mammalian genomes.

It has been proposed that transcription relocalizes (6, 26–29) or interferes (27, 30–34) with cohesin, and that active transcription start sites (TSSs) function as cohesin loading sites (6, 28, 31, 35–37). Induction of genes redistributes cohesin downstream (*i.e*., in the direction of transcription) (26, 29, 38), and cohesin has been observed to accumulate between active convergently transcribed genes, away from putative cohesin loading sites (*e.g*., TSSs, centromeres) (6, 26–29). The emergence of these intergenic “cohesin islands” is particularly prominent in cells in which the cohesin regulators CTCF and Wapl have been depleted (6, 39), but similar accumulation can also occur under physiological conditions, such as senescence (40). In addition, transcription can interfere with cohesin by disrupting its localization at CTCF sites (32), obstructing the growth of cohesin-mediated loops (31, 33), and altering cohesin clustering in 3D (41, 42). Furthermore, single-molecule experiments demonstrated that RNA polymerase (RNAP) can push a passively diffusing cohesin complex along DNA *in vitro* (43). However, it is not known how transcription-driven cohesin relocalization can be reconciled with now well established observations of active loop extrusion by cohesin (3–5, 44, 45), and what patterns of chromatin organization can emerge from transcription-extrusion interactions.

The hypothesis that cohesin loads at active promoters is indirectly supported by ChIP-seq experiments for Nipbl (called Scc2 in yeast), which putatively loads cohesin onto chromatin (46). Nipbl is preferentially detected at the promoters of active genes (6, 35–37, 47–49), which has been interpreted as cohesin preferentially loading at these sites. Similarly, CUT&Tag experiments and computational modeling suggest that promoter-mediated cohesin loading may explain cohesin accumulation at active promoters after mitotic exit (31). However, it is unclear whether preferential loading of cohesin at active TSSs by Nipbl is the primary mechanism accounting for observed cohesin localization and cohesin-mediated genomic contacts.

We sought to unify these observations and determine how transcription modulates loop extrusion to regulate cohesin localization and chromatin organization around genes. We studied cells in which we could alter cohesin abundance, dynamics, and localization. The primary extrusion barriers could be removed by CTCF depletion, and cohesin’s residence time and abundance on chromatin could be increased by Wapl knockout. We found evidence that transcription directly interacts with loop extrusion by cohesin through a “moving barrier” mechanism, similar to how transcription is thought to interfere with condensin in bacteria (50) and yeast (51). Hi-C experiments showed previously unobserved, intricate, cohesin-dependent genomic contact patterns near actively transcribed genes in both wildtype and mutant mouse embryonic fibroblasts (MEFs). In CTCF-Wapl double knockout (DKO) cells (6), genomic contacts were enriched between sites of transcription-driven cohesin localization (cohesin islands). Similar patterns emerged in polymer simulations in which transcribing RNAPs acted as moving barriers by impeding, slowing, and pushing loop-extruding cohesins. Furthermore, the model predicts that cohesin does not load preferentially at promoters and instead accumulates at TSSs due to the barrier function of RNAPs. We tested this prediction by new ChIP-seq experiments with tagged NIPBL. These experiments revealed that the presumed “cohesin loader” Nipbl (46) co-localizes with cohesin, but, unlike in previous reports (6, 35–37, 47–49), Nipbl did not predominantly accumulate at active promoters. Instead, cohesin and Nipbl, as an essential part of the loop-extruding cohesin complex, could accumulate at these sites due to the function of RNAP as a barrier to loop extrusion (9, 30). We propose that RNAP acts as a new type of barrier to loop extrusion that, unlike CTCF, is not stationary in its precise genomic position, but rather, dynamically translocates and relocalizes cohesin along DNA. In this way, loop extrusion could enable translocating RNAPs to maintain contacts with distal regulatory elements, allowing transcriptional activity to shape genomic functional organization.

## Results

### Depletion of Wapl and CTCF shows how transcription governs large-scale genome organization

Since dynamic positioning of cohesin governs global genome organization (52), we investigated whether the relocalization of cohesin by transcription results in large-scale changes to chromatin contacts. We performed high-throughput chromosome conformation capture (Hi-C) in CTCF-Wapl DKO quiescent MEFs and compared it to observations in wild-type (WT), CTCF knockout (KO), Wapl KO, and Smc3 KO cells (**Fig. S1A-B**).

The Hi-C experiments with DKO cells showed new genomic contact patterns generated by cohesin accumulated in “islands” between sites of convergent transcription. We observed new contacts between cohesin islands that appeared as Hi-C “dots” (island-island dots) that bridged distant genomic sites, consistent with the formation of cohesin-mediated chromatin loops (39). While cohesin frequently colocalizes with CTCF in WT cells (6, 11, 12, 17, 53, 54), cohesin islands are not associated with CTCF sites (6) and contacts between CTCF sites are reduced in DKO (**Fig. 1A**). Island-island dots are insulating (comparably to CTCF), and insulation is weakened in Smc3 KO cells (**Fig. S1C-E**). Our findings indicate that cohesins that are relocalized to sites of convergent transcription continue to form large chromatin loops, consistent with ongoing active loop extrusion.

**Figure 1.**
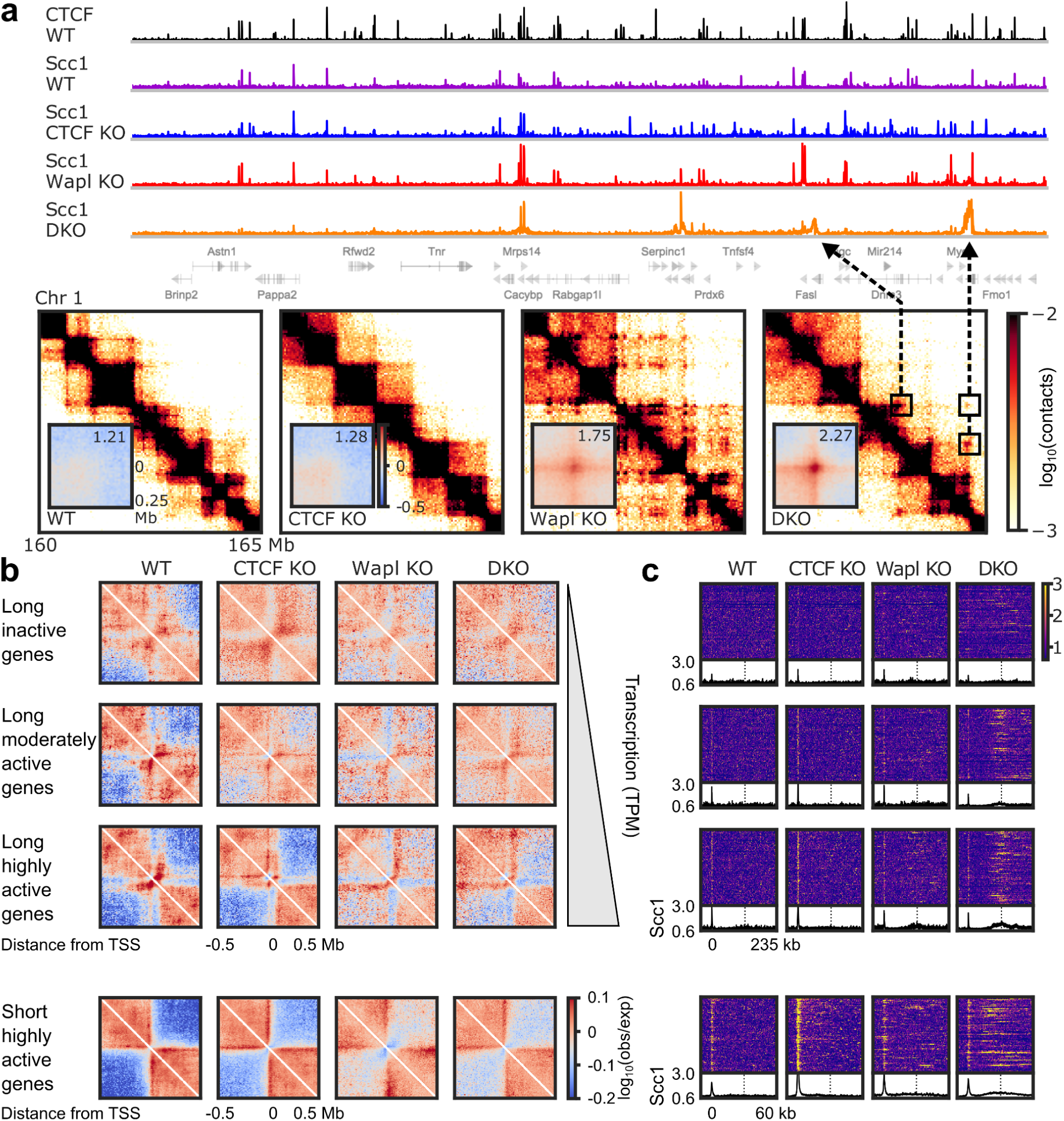
Transcription and cohesin generate characteristic patterns of contacts and cohesin accumulation near active genes. **(A)** *Top*, ChIP-seq tracks for CTCF in WT (black) and Scc1 (cohesin) in WT, CTCF KO, Wapl KO, and DKO cells (purple, blue, red, and orange, respectively) for a 5 Mb region of chromosome 1, with the corresponding gene track below. *Bottom*, Hi-C contact maps for the corresponding region. Boxes in DKO identify examples of island-island dots, with arrows pointing to the corresponding “cohesin islands” in the ChIP-seq tracks. *Insets*, Averages of observed-over-expected contacts (see **Materials and Methods**), centered on island-island dots separated by genomic distances 50 kb < *s* < 350 kb (*n*=1314), plotted with log_10_ color scale. Numbers indicate dot strengths (see **Materials and Methods**). **(B)** Average observed-over-expected Hi-C contact maps centered and oriented on the TSS for long genes (80 kb < *L* < 120 kb), stratified by GRO-seq TPM (top three rows; TPM<0.6, 0.6≤TPM<3.6, and 3.6≤TPM, respectively) and short active genes (bottom row; length 10 kb < *L* < 30 kb; TPM > 3.6). *n*=184, 139, and 176, respectively, for long genes except for DKO, where *n*=123, 233, and 143; *n*=407 for short genes except for DKO, where *n*=592. **(C)** Cohesin (Scc1) ChIP-seq heatmaps and average tracks near long genes stratified by TPM (top three rows) or short active genes (bottom row) oriented and aligned at their TSSs. Heatmaps depict the longest 50% of genes in the group, sorted by decreasing length from top to bottom. Dotted lines in average plots indicate the length of the longest gene in the respective set.

The new “island-island” contacts are clearly distinguishable only in DKO cells (**Fig. 1A**), possibly because they have many more cohesins on chromatin that can be relocalized. CTCF KO abrogates cohesin accumulation at CTCF sites, increasing the quantity of mobile cohesins, and Wapl KO increases cohesin residence time, thus increasing the number of cohesin complexes on chromatin (55) and allowing time for accumulation in islands. Nonetheless, insulation of genomic contacts, and to a lesser degree, cohesin accumulation, also emerge at sites of convergent transcription in WT cells (**Fig. S1D**). Dots, cohesin accumulation, and insulation at cohesin islands depend on transcription. Cohesin accumulation is greater for higher levels of transcription (6), and both dots and insulation at cohesin islands are partially suppressed in DKO by treatment with the transcription elongation inhibitor DRB (**Fig. S1F-G**). Suppression of these features may be only partial due to incomplete inhibition of transcription and that DRB primarily stalls transcription elongation without degrading RNAP (56). Based on these observations, we hypothesize that active transcription may alter genome organization through its effects on loop extrusion by cohesin.

### Cohesin dynamics and transcriptional activity spatially organize chromatin around genes

To directly study how the genome is organized by the interplay of transcription and extrusion, we computed average Hi-C contact maps and Scc1 (cohesin) ChIP-seq tracks centered on transcription start sites (TSSs) of genes, oriented and stratified by transcription activity and gene length (**Figs. 1B-C, Fig. S2,** and **S3A-D**).

In WT and CTCF and Wapl mutants, this revealed that individual genes are insulating, and active genes generate stronger insulation than inactive genes (**Figs. 1B, S2,** and **S3A** and **C**). Contact enrichment and insulation correspond to cohesin accumulation at TSSs (**Figs. 1C** and **S3A-D**). Insulation is abolished in Smc3 KO, while CTCF KO or lack of proximal CTCF only partially weakens insulation (**Figs. 1B, S2,** and **S3E-F**). Insulation is also weakened in Wapl KO and DKO cells (**Fig. 1B**), where increased residence time presumably allows loop-extruding cohesins to traverse the gene and bring regions upstream and downstream of the gene into contact. Thus, active genes are insulating boundaries in both the presence and absence of CTCF, and their effects on local genome organization depend on the dynamics of loop-extruding cohesins.

Near long, active genes, we discovered intricate patterns of genomic contacts and cohesin accumulation, which were modulated by perturbations of cohesin dynamics. Features are better distinguished in long genes than in short genes at least in part due to the resolution of the Hi-C data. Across WT and all mutants (except Smc3 KO; **Fig. S3E**), we observed five major features in these average gene contact maps (**Figs. 1B, 2A, S2,** and **S3A**): 1) insulation (as described above), 2) lines (or “stripes”) of high contact frequency that extend upstream and downstream from the TSS, 3) lines of high contact frequency that extend upstream from the gene body that originate near the 3’ end, 4) high contact frequency within the gene and insulation of the gene, and 5) dots indicating high contact frequency between the 5’ and 3’ ends of the gene. These Hi-C features appeared with sharp ChIP-seq peaks of cohesin accumulation at the TSS, broader cohesin accumulation at the 3’ end of the gene, and a low background level of cohesin within the gene body (**Fig. 1C** and **S3B**). The emergence of lines, dots, and insulation, along with the accumulation of cohesin at the ends of genes, is reminiscent of similar features around CTCF sites and suggests that TSSs and 3’ ends of active genes are barriers to loop extrusion.

**Figure 2.**
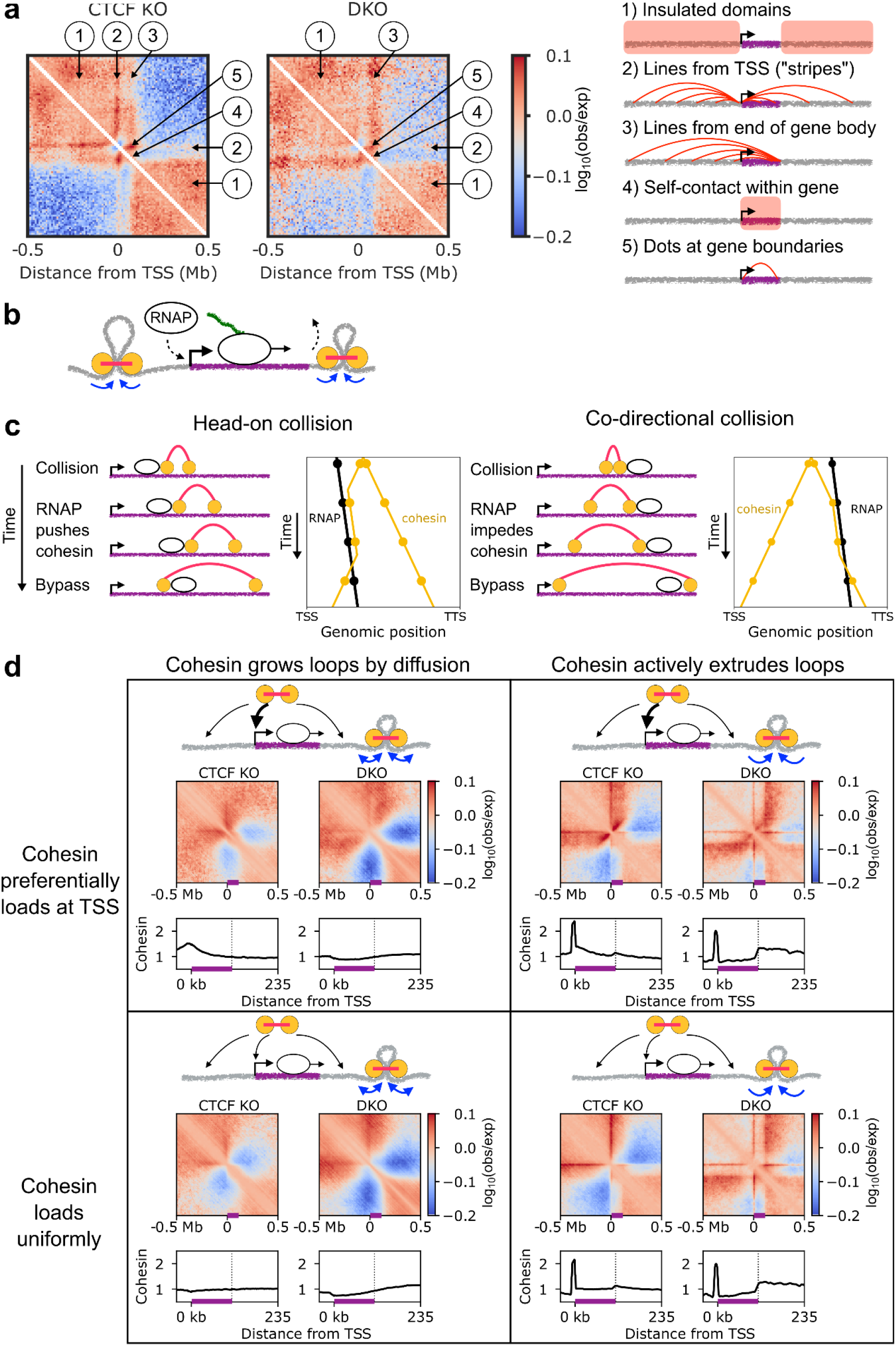
Transcription as a moving barrier for loop extrusion recapitulates major features of genome organization and cohesin accumulation around active genes. **(A)** Observed-over-expected contact maps around long active genes for CTCF KO and DKO with five major features identified and illustrated. **(B)** Schematics of the moving barrier model. Cohesins (yellow and pink) bind to chromatin and extrude loops until unbinding. RNAPs (open ellipses) are loaded at the promoter, translocate through the gene (purple), and are unloaded at the 3’ end. **(C)** Arch diagrams and schematic trajectories illustrating time series of two types of collisions between extruding cohesins and translocating RNAP that may occur in genes in the model. Yellow circles depict the two genomic positions at the base of the extruded loop, bridged by a cohesin. During head-on collisions, RNAP pushes cohesin until the cohesin bypasses the RNAP, the RNAP stops translocating (beyond the 3’ end), or either the RNAP or cohesin unbinds. During co-directional collisions, extrusion by cohesin translocation is slowed by the RNAP barrier moving toward 3’. In both cases, interactions between RNAP and cohesin only alter extrusion on one side of cohesin; collisions do not affect growth of the other side of the extruded loop or RNAP translocation. The trajectory plots show genomic position versus time for RNAP (black) and cohesin’s two sides (yellow). The filled circles indicate the time points and positions corresponding to the illustrations. **(D)** Average observed-over-expected maps and cohesin accumulation tracks near active genes in CTCF KO and DKO simulations. Results shown for simulations with either active extrusion or passive, diffusive loop extrusion, each with either uniform cohesin loading or preferential loading at TSSs. Gene positions are indicated by purple bars on the *x*-axes. *Inset*, illustrations of cohesin loading and translocation.

Consistent with this interpretation, contact patterns are weaker for inactive genes and in cells treated with DRB, especially in DKO (**Fig. 1B, S2,** and **S3C** and **G**). Furthermore, ChIP-seq shows sharp accumulation of RNAP II at the TSS and a smaller, broad accumulation near the 3’ end (**Fig. S3H**), similar to cohesin ChIP-seq (**Figs. 1C** and **S3B**). These observations suggest that RNAPs serve as barriers to loop extrusion, but raise the question of how the spatiotemporal dynamics of RNAP impacts loop extrusion to produce the observed genomic contact patterns.

The lines emanating from 3’ ends of active genes and extending upstream (**Fig. 2A,** feature 3) suggest that 3’ ends are effectively *asymmetric* extrusion barriers. This asymmetry is reminiscent of the lines that emanate from directionally oriented CTCF extrusion barriers (9, 22, 57–59). However, unlike with CTCF barriers, cohesin accumulation is broad (**Figs. 1C** and **S3B**). Together with broad, asymmetric RNAP accumulation at 3’ ends (**Fig. S3H**), these observations suggest that directional RNAP translocation in genes is central to both cohesin accumulation and asymmetric patterns of genomic contacts.

Our findings suggest that transcribing RNAPs are directionally translocating barriers to loop-extruding cohesins, stimulating us to consider a broad class of models for the dynamics and interactions of transcription and loop extrusion.

### The moving barrier model for active loop extrusion can reproduce gene contact maps

#### The moving barrier model

We developed a model to determine how loop-extruding cohesins and their interactions with transcribing RNAPs can generate the major features of contact maps and cohesin accumulation around genes (**Fig. 2B** and **Materials and Methods**). We modeled each cohesin as a two-sided loop-extruding complex that bridges two regions of the chromatin fiber, which are independently and continuously extruded into a chromatin loop (9, 10, 60–62). We considered CTCF KO and DKO scenarios to focus on how transcription affects extrusion without complications from other strong extrusion barriers (*i.e*., CTCF). In DKO simulations, cohesin residence time was increased tenfold and linear density was increased twofold due to Wapl depletion, as suggested by previous experiments (11, 55, 63, 64) and simulations (64, 65).

For extrusion-transcription interactions, we extended the “moving barrier” model for interactions between bacterial condensins and RNAPs (see **Materials and Methods**; (50)). RNAPs load at TSSs, transiently pause, and slowly translocate through the gene (~0.01-0.1 kb/s (66, 67)) and interact with more rapidly translocating loop-extruding cohesins (0.1-1 kb/s (1, 3–5)) (**Fig. 2C**).

When RNAP encounters a cohesin in a head-on collision, it pushes this cohesin along the chromatin fiber in the direction of transcription, shrinking the loop from one side, while the other side of the loop continues to grow at its normal rate (**Fig. 2C**). RNAP pushing cohesin is consistent with the large difference in the stall forces of cohesin (0.1-1 pN; (4, 5, 68)) and RNAP (~10 pN; (66)). Alternatively, when an extruding cohesin progressing toward the 3’ end encounters RNAP, extrusion of that side of the loop continues more slowly behind the slower RNAP, while extrusion continues as normal on the other side of the loop (**Fig. 2C**). In both types of collision, cohesin may stochastically bypass RNAP after a characteristic waiting time, similar to *in vitro* observations of cohesin bypassing obstacles on DNA (69) and predictions for bacterial condensins *in vivo* (50). Thus, in this model, RNAP is a weakly permeable moving barrier to loop extrusion.

We considered four models for cohesin loading and extrusion dynamics (**Fig. 2D**). In our models, cohesin was loaded either uniformly or preferentially at promoters. The latter was suggested by ChIP-seq experiments showing cohesin and Scc2/Nipbl enrichment at TSSs (6, 35–37, 47–49). For each type of cohesin loading, we considered two modes of cohesin extrusion: 1) diffusive growth or shrinking on each side of the extruded loop (70, 71), similar to the earlier hypothesis that RNAPs push passive cohesins to sites of convergent transcription (26, 27) and *in vitro* observations (43), or 2) active, directed loop extrusion of each of the two chromatin strands, as recently observed on DNA *in vitro* (3–5) and suggested by active extrusion models (9, 10, 62).

Using 3D polymer simulations coupled to stochastic 1D transcription and extrusion dynamics (see **Materials and Methods**), we simulated chromosome organization by the moving barrier mechanism with different cohesin loading scenarios (loading uniformly or preferentially at TSS), loop extrusion activities (active or passive), and cohesin-RNAP bypassing times.

#### Loop extrusion with moving barriers generates experimentally observed genomic contact patterns

The four models with different cohesin loading scenarios and loop extrusion mechanisms produced different contact maps around active genes, allowing us to select the class of models that best matches the experiments.

In models with diffusively, rather than actively, extruding cohesins, active genes alter cohesin accumulation patterns and the spatial organization of the genome, but the simulations lacked prominent features observed via Hi-C and ChIP-seq, irrespective of the loading scenario, diffusion coefficient, and cohesin-RNAP bypassing time (**Figs. 2D** and **S4)**. Cohesin accumulation, where it occurred, was weak and broad because diffusive cohesins do not remain localized after encountering an extrusion barrier. This resulted in weak, poorly defined features in simulation contact maps. We conclude that diffusively extruding cohesins do not reproduce the experimental observations around active genes, even when they are pushed by RNAPs.

In contrast, simulations with cohesins that actively extrude loops produced genome contact maps and cohesin accumulation patterns with well defined features (**Figs. 2D** and **S5**). Active extrusion with a low, but nonzero, rate of cohesin-RNAP bypassing (~1 event per cohesin lifetime, *i.e*., cohesin follows or is pushed by RNAP for ~100 s) gave the best agreement with experiments. Both with and without preferential loading at the TSS, cohesin sharply accumulated at TSSs and more broadly accumulated at 3’ ends of genes, similar to experimental observations. TSS accumulation occurred because RNAPs that occupied the TSS prior to initiation acted as barriers to extrusion (**Fig. S6** and **S7**). Cohesin accumulated near 3’ by two mechanisms: 1) RNAPs paused, but still bound, at the 3’ end after transcription termination act as barriers and 2) translocating RNAPs that encounter extruding cohesins head-on push the cohesins back toward 3’ ends and slow down extrusion by trailing cohesins (**Fig. 2C** and **S8**). Consistent with these mechanisms, cohesin accumulation at 3’ gene ends was enhanced in DKO simulations due to their longer residence time. In both CTCF KO and DKO simulations, cohesin accumulation resulted in insulation (feature 1), lines emanating from the TSS (feature 2), and lines running upstream from 3’ ends (feature 3). Consistent with the experimental observations, Hi-C lines from 3’ were particularly thick in DKO simulations. We also observed enrichment of contacts within the gene and insulation of the gene (feature 4), as well as dots for contacts between gene ends (feature 5). Therefore, the simple moving barrier model with active extrusion reproduced the major features of active gene organization remarkably well.

Our model also allows us to differentiate between two previously proposed (9, 28, 35) modes of cohesin loading. Extrusion with uniform cohesin loading reproduced the experimental Hi-C maps better than models with a strong preference for loading at an active promoter (**Fig. 2D**). In contrast, simulations with targeted loading had an additional strong feature that is not present in the experiments: diagonal lines that emanate from the TSS, perpendicular to the main diagonal. These lines of enriched contacts formed because cohesins loaded at the TSS brought chromatin on both sides of the TSS together as they progressively extruded loops, reminiscent of patterns emerging when bacterial condensins are loaded at *parS* sites (72–74). This observation suggests that strong preferential loading of cohesin at all promoters is inconsistent with genome organization around active genes.

We next investigated whether the active translocation by RNAP is necessary to generate the genomic contact patterns observed in experiments. We performed simulations with stationary RNAP barriers distributed randomly throughout the genes (see **Supplemental Methods**). We observed that the characteristic features of transcription-extrusion interactions are present in these simulations, but they are weakened compared to simulations with translocating RNAP barriers (**Fig. S9**). These results are reminiscent of the subtle effects observed in experiments with DRB treatment (**Fig. S3G**), which stalls RNAP (56). Furthermore, these simulations demonstrate that the active shuttling of extruding cohesins by RNAP is an important part of the mechanism generating genomic contact patterns around active genes.

The moving barrier model also makes several testable predictions about genome contact patterns. First, it reproduces the experimental observations of cohesin islands and island-island dots at sites of convergent transcription in DKO cells (**Fig. S10**). The simulations additionally predict that the TSSs of divergently oriented genes form contacts (dots) in both CTCF KO and DKO simulations (**Fig. S10**). This is consistent with the idea that TSSs occupied by RNAPs are barriers to loop extrusion that can accumulate cohesin. Since cohesin also accumulates at the 3’ ends (**Figs. 1C** and **2D**), the model predicts an enrichment of contacts between two consecutive ends of active genes, regardless of orientation. The simulations further suggest that cohesin is uniformly loaded on chromatin, without a strong preference for loading at the promoter (**Fig. 2D**). We next tested these two predictions by new ChIP-seq experiments and analysis of Hi-C data.

### Transcription generates genomic contacts between gene ends

To test the prediction that nearby ends of active genes are barriers to extrusion with enriched genomic contacts, we computed average contact maps for contacts between gene ends for pairs of active genes of various orientations. Across WT and all mutants with cohesin, we observed dots of high contact frequency between proximal TSSs (**Fig. 3A**), and contacts between TSSs of adjacent active genes can be observed for all gene pairs regardless of their orientations (**Fig. S11A**). As predicted, 3’ ends of genes can also act as extrusion barriers that enhance contacts between genes (**Fig. S11A**). In each case, contacts are weakened when cohesin residence time is increased by Wapl depletion, presumably because the longer residence increases the probability of cohesin translocating through permeable RNAP barriers and the gene. Contact enrichment depends on transcription, as dots are not observed for inactive genes (**Fig. S11B**). These results demonstrate that cohesins generate specific genomic contacts in response to the cell’s transcriptional activity.

**Figure 3.**
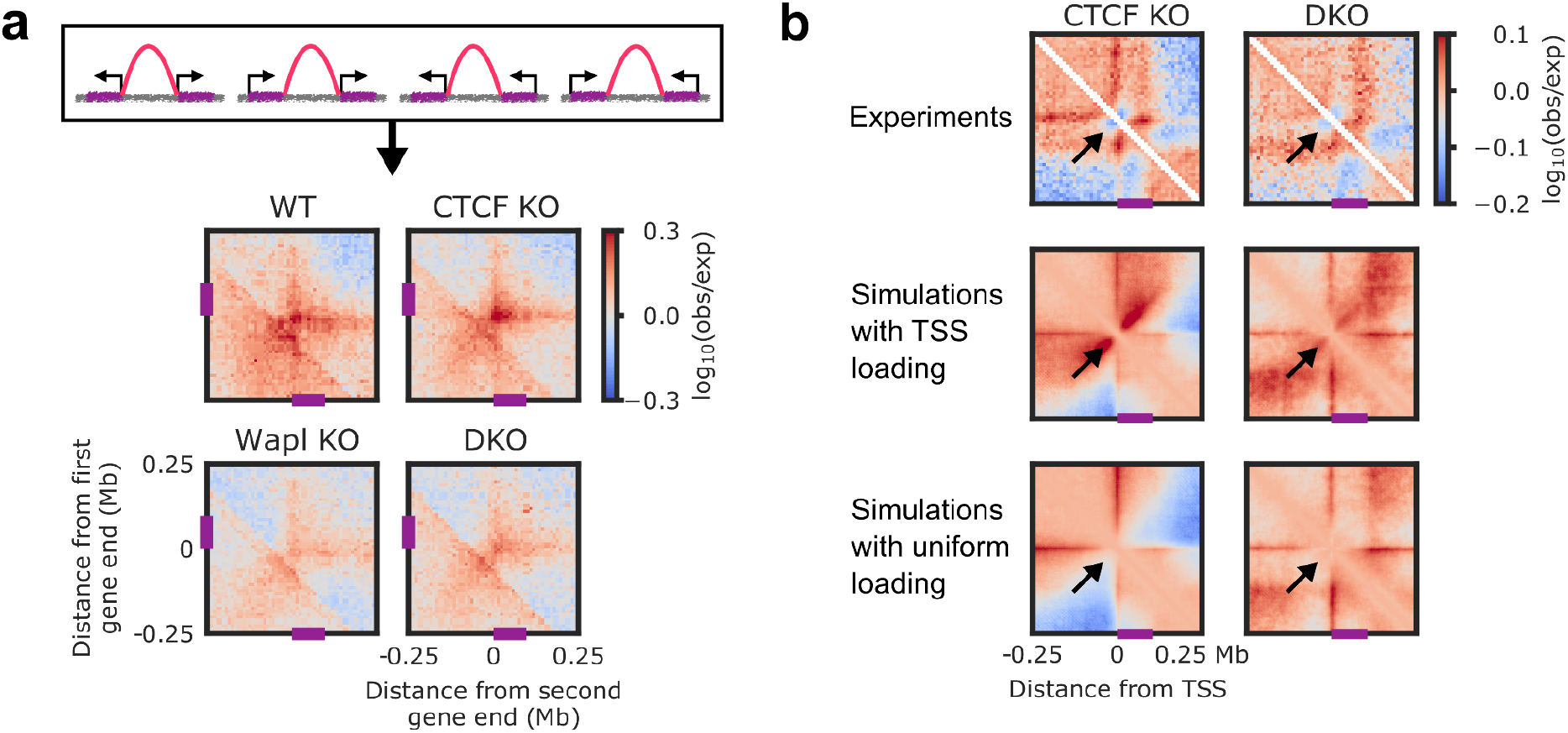
Predictions of the moving barrier model. **(A)** *Left*, Drawing showing contacts between ends of nearby genes in four pairs of orientations. *Right*, Average observed-over-expected maps centered on contacts between nearest ends of active (TPM > 2) genes separated by 50 kb < *s* < 350 kb. At least one gene in each pair is not near a CTCF site (see **Materials and Methods**). **(B)** Zoomed-in views of average observed-over-expected maps for CTCF KO and DKO in experiments and simulations with active, directed loop extrusion, with and without preferential cohesin loading at TSSs. Arrows indicate the presence or lack of an extra line of enriched genomic contacts characteristic of cohesin loading at the TSS. Gene positions are shown by purple bars on the axes.

### The ‘cohesin loader’ NIPBL is not predominantly enriched at promoters

It is widely held that cohesin is loaded preferentially at promoters of active genes, but our moving barrier model, on the contrary, predicts that uniform cohesin loading along the chromatin fiber better recapitulates the Hi-C data (**Figs. 2D** and **3B**). The hypothesis that cohesin loading occurs at TSSs is largely based on the notion that cohesin is loaded onto DNA by NIPBL (46) and that in ChIP-seq experiments NIPBL antibodies preferentially detect TSSs (6, 35–37, 47–49). However, there is no direct evidence that cohesin is loaded onto DNA at sites at which NIPBL ChIP-seq signals have been detected, and alternatively, these sites could represent the presence of extruding cohesin, which also contains NIPBL (3, 4). Furthermore, it is unclear how specific these NIPBL signals are since active TSSs have been identified as ‘hyper-ChIP-able’ regions that some antibodies recruited even in the absence of their antigen (75–77), and negative controls have not been reported for NIPBL ChIP-seq experiments. We therefore reexamined the enrichment and localization of NIPBL throughout the genome.

For this purpose, we generated HeLa cell lines in which all NIPBL or MAU2 alleles were modified with a hemagglutinin tag (HA) and FK506 binding protein 12-F36V (FKBP12^F36V^) (**Fig. S12A**). The HA tag can be specifically recognized in ChIP-seq experiments and FKBP12^F36V^ can be used to induce degradation of the resulting fusion proteins (78) to perform negative control experiments (**Fig. S12B**). The resulting HA-FKBP12^F36V^-NIPBL (HA-NIPBL) and MAU2-FKBP12^F36V^-HA (MAU2-HA) fusion proteins were functional since they supported formation of vermicelli, axial chromosomal sites at which cohesin accumulates in Wapl-depleted cells (55) in a manner that depends on NIPBL (63, 79)) (**Fig. S12C-D**).

HA-NIPBL and MAU2-HA ChIP-seq experiments identified small numbers of peaks, most of which disappeared upon dTAG-induced degradation (**Figs. 4A-B** and **S12E**). Of these ‘high-confidence’ NIPBL-MAU2 peaks, which we used as a reference set (see **Materials and Methods**), 86% overlapped with cohesin peaks, but only 16.7% were located at TSSs (**Fig. 4C**). Moreover, nearly all the latter NIPBL-MAU2 peaks were located at TSSs at which cohesin was also enriched (**Fig. 4B-C**).

**Figure 4.**
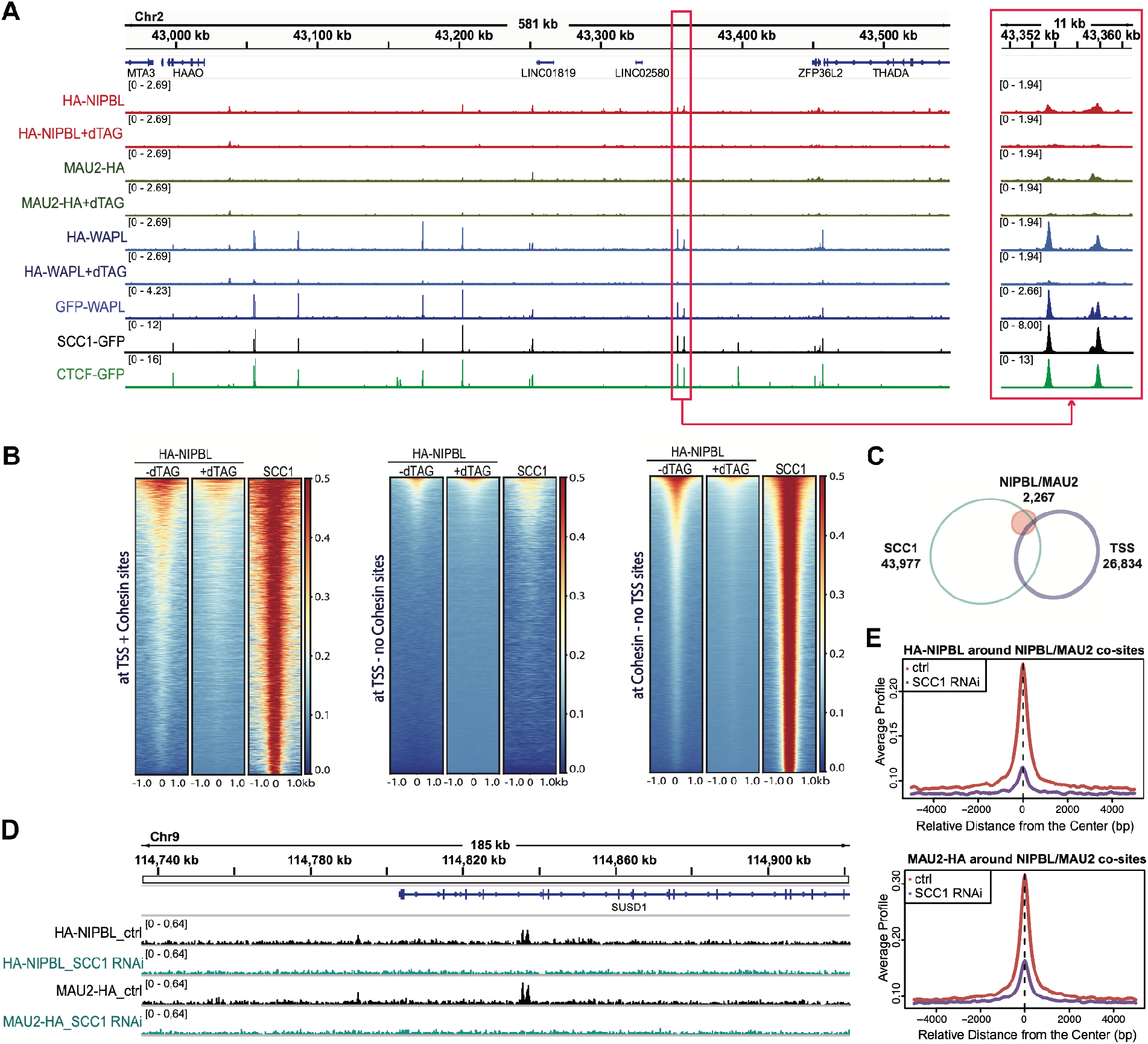
NIPBL and MAU2 colocalize predominantly with cohesin but not TSS throughout the genome. **(A)** ChIP-Seq profiles of HA-NIPBL (-/+dTAG), MAU2-HA (-/+dTAG), HA-WAPL (-/+dTAG), GFP-WAPL, SCC1-GFP and CTCF-GFP along an exemplary 581 kb region of chromosome 2, illustrating the typical distribution and colocalization of sequencing read pileups. Genes within this region are depicted above. The red rectangle on the left indicates one region of interest and a zoom-in view is shown on the right. **(B)** Heatmaps of HA-NIPBL (-/+dTAG) and SCC1 ChIP-Seq at TSSs with cohesin, TSSs without cohesin, and cohesin sites not at TSSs. **(C)** Area-proportional threefold eulerAPE Venn diagram illustrating overlap between NIPBL/MAU2 co-sites, SCC1, and TSS. **(D)** Enrichment profiles of HA-NIPBL (-/+ SCC1 RNAi) and MAU2-HA (-/+ SCC1 RNAi) along an exemplary 185 kb region of chromosome 9, illustrating typical distribution and colocalization of sequencing read pileups. Genes within this region are depicted above. **(E)** Average signal profiles of HA-NIPBL (-/+ SCC1 RNAi) or MAU2 (-/+ SCC1 RNAi) around NIPBL/MAU2 co-sites.

Since NIPBL and MAU2 co-localize with cohesin, we hypothesized that NIPBL-MAU2 complexes might be recruited to these sites by binding to cohesin. To test this possibility, we analyzed whether HA-NIPBL and MAU2-HA ChIP-seq peaks depend on cohesin by depleting SCC1 via RNAi (**Figs. 4D-E** and **S12F**). Both HA-NIPBL and MAU2-HA ChIP-seq peaks were greatly reduced or undetectable after depletion of SCC1. These experiments suggest that NIPBL-MAU2 complexes are not enriched at TSSs unless these are also occupied by cohesin; rather, NIPBL-MAU2 colocalizes with a subset of cohesin complexes on chromatin.

We also tested the specificity of two NIPBL antibodies that have been used in previous studies by performing ChIP-seq experiments with HeLa cells from which NIPBL had been depleted or not. One of these antibodies is “133M” (37). The other one is available from Bethyl Laboratories (“Bethyl”). In these experiments, we used unmodified HeLa cells and depleted NIPBL by RNAi to rule out the possibility that tagging NIPBL with HA and FKBP12^F36V^ would alter recognition of NIPBL by these antibodies. We controlled the depletion of NIPBL by immunoblot analysis of its binding partner MAU2 (**Fig. S13A**), because NIPBL degradation leads to depletion of MAU2, which can be analyzed by immunoblotting more reliably than the 316 kDa NIPBL protein (80).

In our experiments, 133M antibodies identified 11,001 peaks, of which 7,774 were located at TSSs but only 4,547 overlapped with cohesin (**Fig. S13B-C** and **S14A-C**), similar to previous observations (37). However, after NIPBL depletion by RNAi, 9,596 (87%) ChIP-seq peaks remained (**Figs. S13B-C** and **S14**), suggesting that most of these peaks did not depend on NIPBL. The Bethyl antibodies identified 6,587 peaks, of which only 2,445 were located at TSSs but 5,008 overlapped with cohesin (**Figs. S13B-C** and **S14D-F**). Of the peaks detectable with the Bethyl antibodies, only 2,093 (32%) remained after NIPBL RNAi. Furthermore, peaks detected by the Bethyl antibodies covered a higher fraction (54%) of NIPBL/MAU2 peaks detected by tagging these proteins with HA than those detected by the 133M antibodies (23%) (**Fig. S14**). These results indicate that only some peaks detected by 133M but most peaks detected by Bethyl depend on NIPBL, and the majority of the specific peaks overlap with cohesin.

Together with our simulations (**Fig. 3B**), these findings suggest that NIPBL does not primarily accumulate at TSSs and that cohesin complexes are not preferentially loaded onto chromatin at these sites. Instead, cohesin accumulation at TSSs could occur due to the barrier function of TSSs, and in turn, NIPBL could colocalize with cohesin at sites where loop extrusion is impeded.

## Discussion

It has been hypothesized that RNAPs push cohesin complexes that have entrapped DNA within their ring structures, displacing cohesins from their apparent loading sites at gene promoters (35) to the 3’ ends of genes (6, 26–29). Indeed, previous single-molecule experiments demonstrated that a transcribing RNAP could push a passively diffusing cohesin complex along DNA *in vitro* (43). However, it is now known that cohesin can translocate by actively extruding DNA loops (3–5). Cohesin can do so without topologically encircling DNA (3), and furthermore, it can bypass large obstacles on DNA (69). The mechanism introduced here can reconcile these new developments with older observations of the effects of RNAP on cohesin.

Our experiments and simulations indicate that RNAP acts as a “moving barrier” to loop-extruding cohesin. RNAP in a head-on collision with cohesin can push cohesin toward a gene’s 3’ end as cohesin continues to extrude at the other end of its loop. Alternatively, RNAP can slow extrusion by cohesin trailing the RNAP, while cohesin can continue to rapidly extrude the other side of the loop (**Fig. 2A-B**). Our simulations suggest that extrusion should be at least 3-5 times faster than transcription to obtain the experimentally observed genomic contact patterns (**Fig. S8**). RNAPs accumulated at TSSs and transcription termination sites also act as extrusion boundaries that enrich contacts between nearby gene ends (**Figs. 3A** and **S11**), and stationary RNAPs within the gene can also provide some degree of organization (**Figs. S3G** and **S9**). These findings generalize the bacterial moving barrier model (50) to eukaryotic cells and provide a detailed account of how transcription interacts with extrusion (30, 31, 33, 51) to locally modulate genome organization by stopping, hindering, and relocalizing cohesins.

The effects of transcription on extrusion are most clearly visible in CTCF-Wapl DKO cells, which in turn provides insights relevant in WT cells. In DKO, the strong interfering signal from CTCF barriers is suppressed and cohesin is long-lived, allowing it to be pushed or impeded over long distances by RNAP (up to ~10 kb in WT versus ~100 kb DKO in simulations). These differences in cohesin dynamics strengthen some features in DKO, particularly lines running upstream from the 3’ end of the gene (feature 3 in **Results** and **Fig. 2A**) and 3’ cohesin accumulation (**Fig. 1C**), supporting the conclusion that RNAP can push extruding cohesins. Other features, such as insulation (feature 1) are weakened, indicating that extruding cohesin can traverse the gene and bypass RNAPs during its longer lifetime. These effects are clearest in long genes, due to both Hi-C resolution and the larger total probability of cohesin encountering RNAP. Altogether, the differences between WT and DKO demonstrate how transcription-extrusion interactions can manifest differently according to the dynamics of loop extrusion by cohesin and can modulate transcription-driven chromatin organization.

Accordingly, through moving barrier interactions, transcription can differentially and locally shape genome architecture in a variety of physiological scenarios. Normal transcriptional responses or excess readthrough can disrupt cohesin- and CTCF-mediated looping within genes and near 3’ ends in human cells (32, 34). Through the moving barrier mechanism, transcription can also *enrich* contacts, as it does between genes in mouse cells (**Fig. 3A** and (21, 81)) or between sites of convergent transcription in both yeast (33, 82) and mammalian cells (**Fig. 1A** and (39, 40)), similar to our observations in DKO cells (**Fig. 1A**). Furthermore, in mammalian cells, TSSs and 3’ ends of active genes insulate genomic contacts (21, 30, 81, 83–87) by acting as extrusion boundaries (**Figs. 1B-C**, **2, 3A**, **S11**, and **S15**; (30)). Thus, transcription-extrusion interactions appear to locally organize chromatin in a variety of physiological scenarios.

Functionally, RNAP barriers could regulate genes through their effects on loop extrusion. Pausing of extrusion at TSSs would allow cohesin to linearly scan chromatin for proximal enhancers near other boundaries, such as CTCFs, to bring them into contact with the TSS (88). Paused RNAPs or RNAPs initiating a lower level of transcription may stop extrusion on one side of cohesin, while allowing the other side to scan to an enhancer, which in turn could trigger a higher level of expression. Subsequently, cohesin may track the transcribing RNAP through the gene, maintaining continuous contact between the enhancer and the transcription complex, as previously observed (89). Further, cohesin-mediated contacts between nearby active promoters (**Fig. 3A** and (21, 81)) may mediate mutual regulation, possibly facilitating the spreading of histone marks and transcription factors. Linear scanning by cohesin in gene regulation would also be consistent with cohesin’s proposed role in other contexts, including V(D)J recombination (23, 90–93), alternative protocadherin choice (24), and double strand break repair (94, 95). Furthermore, since moving RNAP extrusion barriers are transcription-dependent, chromosomal interactions could be rapidly modulated in a locus-specific manner. For instance, histone marks or transcription factors could regulate genomic contacts by locally activating transcription, thus modulating functional interactions in *cis* via extrusion. Similarly, even non-protein-coding genes could have this effect through, for example, transcription of non-coding RNAs and eRNAs (96, 97). Thus, while cohesin may only moderately alter global transcription (6, 8–11, 98), cohesin-RNAP interactions could impact transcription of specific genes that depend on the recruitment of nearby *cis* regulatory elements. Therefore, in contrast to static barriers like CTCFs, moving RNAP barriers can dynamically regulate loop extrusion and functional interactions.

Our modeling predicts that cohesin is not preferentially loaded at active promoters (**Figs. 2D** and **3B**), in contrast to previous proposals (6, 31, 35–37, 48, 49). Consistent with our prediction, our new ChIP-seq experiments (**Fig. 4**) show that the enrichment of the “cohesin loader” NIPBL at TSSs may have been, at least in part, an artifact, possibly because active TSSs are ‘hyper-chippable,’ especially when the antibody has a limited specificity (75–77). Furthermore, we found that NIPBL occupancy depends on the presence of cohesin (**Fig. 4D-E**), consistent with the requirement of NIPBL for *in vitro* loop extrusion (3, 4) and *in vivo* loop lengthening (63, 79) (**Fig. S12C-D**) by cohesin. Peaks of NIPBL and cohesin accumulation may reflect stopping the translocation of the entire extruding complex, rather than loading. In fact, we demonstrated that loading would leave a distinct diagonal pattern not observed at TSSs (**Fig. 3B**). This further suggests that NIPBL may serve as an extrusion processivity factor for cohesin rather than only as a loading factor (3, 52).

Even though RNAP relocalizes and slows cohesin in our model, cohesin that is pushed or impeded by RNAP can bypass RNAP approximately once per 100 s (which is also the simulated WT cohesin lifetime). This suggests that cohesin may translocate with RNAP for distances of order 10 kb in WT and 100 kb in Wapl KO cells before bypassing occurs. The ability of cohesin to bypass RNAP is consistent with experiments indicating that cohesin does not topologically enclose DNA while it extrudes loops (3, 69). However, our model’s bypassing time is 10-fold longer than predicted for bacterial condensins bypassing RNAPs (50) and measured for SMC complexes bypassing obstacles on DNA *in vitro* (69). This discrepancy suggests differences between loop extrusion on nucleosomal fibers versus DNA or that cohesin may have some affinity for RNAP. The former could be due to steric interactions imposed by nucleosomes and/or large nascent RNA molecules trailing the RNAP, while the latter could facilitate the linear scanning processes described above. However, much like the mechanism of extrusion itself, the molecular mechanisms of interactions with RNAP and bypassing remains unclear.

Our results indicate that RNAP belongs to a growing list of elements that dynamically structure the genome by acting as barriers to loop extrusion. However, while boundaries such as CTCF sites are stationary (6, 8–12), RNAPs are mobile and can be dynamically controlled by transcriptional regulators. Together with emerging evidence that extrusion might also be obstructed by other mobile complexes such as replication machinery (25, 33), this suggests that in addition to *structural* functions, cohesin has important *dynamic* functions in the spatiotemporal organization of the genome.

## Materials and Methods

Descriptions of HeLa cell lines, antibodies and reagents, whole cell extract, chromatin fractionation, immunofluorescence microscopy, RNAi, additional details of ChIP-seq, simulations with stationary RNAP, and implementation of 3D polymer dynamics in simulations can be found in the **Supplemental Methods**.

### Hi-C protocol for MEFs

Hi-C was performed as described previously (99). Briefly, 30×106 cells were cross-linked in 2% formaldehyde for 10 minutes and quenched with ice-cold glycine (0.125 M final concentration). Cells were snap-frozen and stored at −80°C before cell lysis. Cells were lysed for 30 min in ice cold lysis buffer (10 mM Tris-HCl, pH 7.5, 10 mM NaCl, 5 mM MgCl2, 0.1 mM EGTA, and 0.2% NP-40) in the presence of protease inhibitors. Chromatin was solubilized in 0.6% SDS at 37°C for 2h minutes, quenched by 3.3% Triton X-100. Chromatin was digested with 400 units of HindIII overnight at 37°C. Fill-in of digested overhangs by DNA polymerase I, large Klenow fragment in the presence of 250 nM biotin-14-dATP for 90min was performed prior to 1% SDS based enzyme inactivation and dilute ligation with T4DNA ligase for 4 hours at 16°C. Cross-links of ligated chromatin were reversed overnight by 1% proteinase K incubation at 65°C. DNA was isolated with 1:1 phenol:chloroform, followed by 30 minutes of RNase A incubation. Biotin was removed from unligated ends by incubation with 15 units of T4 DNA polymerase. DNA was sheared using an E220 evolution sonicator (Covaris, E220) and size selected to 150-350 bps by using AMPure XP beads. After end repair in a mixture of T4 polynucleotide kinase, T4 DNA polymerase and DNA polymerase I, large (Klenow) fragment at room temperature for 30 minutes, dATP was added to blunted ends polymerase I, large fragment (Klenow 3’ → 5’ exo-) at 37°C for 30 minutes. Biotinylated DNA was collected by incubation in the presence of 10 μl of streptavidin coated myOne C1 beads and Illumina paired-end adapters were added by ligation with T4 DNA ligase for 2 hours at room temperature. A PCR titration (primers PE1.0 and PE2.0) was performed prior to a production PCR to determine the minimal number of PCR cycles needed to generate a Hi-C library. Primers were separated from the library using AMPure XP size selection prior to 50 bp paired-end sequencing (HiSeqv4, Illumina).

### Hi-C mapping and analysis

Hi-C data were mapped to 1 kb resolution using the mm9 genome assembly and distiller pipeline (https://github.com/mirnylab/distiller-nf; version 0.0.3 for all datasets except for DRB and DRB release, which used version 0.3.1). For wildtype and each mutant, >430 million total reads were recorded and >320 million reads were mapped. The mapped data were converted to cooler files (100) and balanced by iterative correction as described previously (101). Contact probability scalings, *P_c_*(*s*), and insulation were computed using cooltools (https://github.com/mirnylab/cooltools; (100)). Pile ups were computed from Python scripts by collecting snippets of maps (“observed”) around sites of interest (such as ends of genes, CTCF sites, or island-island contacts), normalizing each diagonal by the value of the scaling (“expected”) at that diagonal, and averaging “observed-over-expected” values across the collected snippets (https://github.com/mirnylab/moving-barriers-paper). To select Hi-C regions around genes based on transcription levels and gene length, we combined gene annotations for genes with a known transcription status from GENCODE (https://www.gencodegenes.org/) with previously reported GRO-seq for MEFs ((6); GEO accession number GSE76303). Unless noted, we considered only genes isolated from other genes by at least 10 kb. Dot strengths are computed by summing observed-over-expected within a 50 kb of the dot and dividing by the background, taken to be the mean number of contacts in two windows of the same size centered 150 kb upstream and downstream of the dot. For analyzing genomic loci, such as genes, that are “away” from CTCF sites, unless noted, we excluded sites within 5 kb of the top 50%, by motif score, of identified CTCF sites.

### Calibrated ChIP followed by next-generation sequencing

ChIP was performed as described previously (12). Before crosslinking, 10 million HeLa cells were spiked in with 5% MEFs cells (except for RNA polymerase II S2 ChIP, which used only 10 million MEFs cells, without spike-in). Cells were crosslinked with 1% formaldehyde at room temperature for 10 min and subsequently quenched with 125 mM glycine for 5 min. Cells were washed with PBS and then lysed in lysis buffer (50 mM Tris-HCl pH 8.0, 10 mM EDTA pH 8.0, 1% SDS, protease inhibitors) on ice for 10 min. DNA was sonicated by 6 cycles (30 sec on/off) using Biorupter. 10 volumes of dilution buffer (20 mM Tris-HCl pH 8.0, 2 mM EDTA pH 8.0, 1% Triton X-100, 150 mM NaCl, 1 mM PMSF) was added to the lysate, and followed by pre-clear with 100 μl Affi-Prep Protein A beads at 4°C. Immunoprecipitation was performed with rabbit IgG or antibody overnight, followed by 3-hour incubation with Affi-Prep Protein A beads. Beads were washed twice with Wash buffer 1 (20 mM Tris-HCl pH 8.0, 2 mM EDTA pH 8.0, 1% Triton X-100, 150 mM NaCl, 0.1% SDS, 1 mM PMSF), twice with Wash buffer 2 (20 mM Tris-HCl pH 8.0, 2 mM EDTA pH 8.0, 1% Triton X-100, 500 mM NaCl, 0.1% SDS, 1 mM PMSF), twice with Wash buffer 3 (10 mM Tris-HCl pH 8.0, 2 mM EDTA pH 8.0, 250 mM LiCl, 0.5% NP-40, 0.5% deoxycholate), twice with TE buffer (10 mM Tris-HCl pH 8.0, 1 mM EDTA pH 8.0), and eluted twice with 200 μl elution buffer (25 mM Tris-HCl pH 7.5, 5 mM EDTA pH 8.0, 0.5% SDS) by shaking at 65°C for 20 min. The eluates were treated with RNase-A at 37°C for 1 hour and proteinase K at 65°C overnight. Addition of 1 μl glycogen (20 mg/ml) and 1/10th volume sodium acetate (3 M, pH 5.2) was followed by extraction with phenol/chloroform/isoamyl alcohol (25:24:1), precipitation with ethanol. DNA was re-suspended in 100 l of H2O, and ChIP efficiency was quantified by quantitative PCR (qPCR). The DNA samples were submitted for library preparation and Illumina deep sequencing to the Campus Science Support Facility. Depth of sequencing was 50 bp. For Pol II ChIP-seq, >46 million total reads were recorded and over >39 million were uniquely mapped for each condition. For NIPBL, HA-NIPBL, HA-WAPL, and MAU2-HA ChIP-seq, >71 million total reads and >56 million uniquely mapped reads were recorded for each condition.

### Polymer simulations with loop extrusion

Polymer simulations with loop extrusion were performed using OpenMM (102, 103) and the openmm-polymer library (https://github.com/mirnylab/openmm-polymer-legacy), as described previously (9, 62, 104). The simulation code implementing the moving barrier model with these packages is freely and publicly available (https://github.com/mirnylab/moving-barriers-paper). Loop extrusion dynamics with RNAP moving barriers are computed through the 1D model described below. Genomic positions of loop extruders as a function of time determine which monomeric subunits of the polymer are bridged at any particular instant in 3D simulations.

#### Computation of loop extrusion dynamics

##### Cohesin dynamics

The chromosome is modeled as a 1D array of *L*=10^4^ genomic (lattice) sites, each of which represents 1 kb of chromatin. Cohesin complexes are modeled as loop-extruding factors (LEFs) with two linked components. Each component of a cohesin complex occupies a distinct, single lattice space. A LEF is loaded onto a pair of adjacent lattice sites that is not occupied by another LEF or RNAP. At each subsequent timestep, each component of the LEF may translocate to an unoccupied adjacent site with a probability determined by the type of extrusion dynamics simulated. Each of the two components in an individual LEF translocates (or not) independently of the other. For active, directed extrusion, each LEF component translocates away from its initial loading site (growing the loop) onto the next adjacent site with probability *v*=1, provided that the new lattice site is unoccupied; LEF components stop when they encounter another LEF component. For passive, diffusive extrusion, a LEF component may translocate in either direction along the chromosome lattice with equal probability, again provided that the new lattice site is unoccupied. Each component translocates in a single direction during each timestep, with probability *v*=0.5 per possible move. In both cases, extrusion proceeds in this manner (modulated by interactions with RNAPs; see below) until unbinding from the chromosome, which occurs with probability 1/*τ*, where *τ* is the mean residence time. Upon unbinding, the LEF is instantaneously reloaded to another pair of lattice sites. Loading biases are implemented by increasing the probability of binding to the TSS to 100-fold that of other lattice sites.

For active extrusion, *τ*=100 for CTCF KO and *τ*=1000 for DKO simulations, giving mean LEF processivities of *λ*=200 kb and 2000 kb, respectively, similar to previous studies (62, 65). In passive extrusion simulations, standard parameters were *τ*=5000 or 50000 for CTCF KO or DKO, respectively. Setting the extrusion speed to *v*=1 kb/s, sets 1 timestep = 1 s in directed extrusion simulations. In simulations with passive extrusion, rescaling the lifetimes to match the directed case sets 1 timestep = 0.02 s and the LEF diffusion coefficient to *D*=12.5 kb^2^/s, consistent with previous simulations (62) and experimental measurements (43, 105, 106). For simulations with slow diffusive extrusion, *τ*=100 for CTCF KO and *τ*=1000 for DKO, and *v*=0.07 kb/s (*D*=0.005 kb^2^/s).

Simulations are performed with *N*=*L/d*=50 LEFs in CTCF KO and *N*=100 LEFs in DKO (65), giving a mean separation *d*=200 kb and 100 kb, respectively.

##### RNA polymerase dynamics

RNAP barriers were incorporated by adapting the moving barrier model for bacteria (50). To simulate contact patterns near genes, we simulated chromosomes with 9 genes, with TSSs located at *s*=950, 1950, 2950, …, 8950 kb and transcription termination sites (TTSs) located 110 lattice sites (110 kb) downstream. For simulations of genes in convergent orientations, convergently oriented gene pairs had TSSs at (*s*_1_, *s*_2_)= (840 kb, 1160 kb), (1640 kb, 1960 kb), (2440 kb, 2760 kb), … (8840 kb, 9160 kb); thus, the sites of convergent transcription used for pile ups were located at *s*=1000, 1800, 2600, …, 9000 kb. RNAP may be loaded onto an unoccupied TSS with at rate *k*_load_. It remains paused at the TSS until it unpauses at rate *k*_unpause_. After unpausing, the RNAP stochastically translocates by at most one lattice site per timestep toward the TTS at rate *v_p_*, provided the site is unoccupied by another RNAP. After passing the TTS, the RNAP stalls at rate *k*_stall_, and subsequently unbinds at rate *k*_unbind_. Only one RNAP may occupy a single lattice site.

In simulations with directed extrusion, *k*_load_=0.001, *k*_unpause_=0.002, *v_p_*=0.1, *k*_stall_=0.001, and *k*_unbind_=0.002, unless noted. These correspond to *k*_load_=0.001 s^-1^, *k*_unpause_=0.002 s^-1^, *v_p_*=0.1 kb/s (66, 67), *k*_stall_=0.001 s^-1^, *k*_unbind_=0.002 s^-1^. Times (and thus, rates) are rescaled in passive extrusion simulations according to the cohesin residence time, so there, *k*_load_=2·10^-5^, *k*_unpause_=4·10^-5^, *v_p_*=0.002, *k*_stall_=2·10^-5^, and *k*_unbind_=4·10^-5^. Simulations with different *k*_unpause_ and *v_p_* are shown in **Figs. S6** and **S8**.

##### Cohesin-polymerase interactions

When a RNAP and a LEF arrive at adjacent lattice sites and the RNAP is translocating toward the LEF, they are in a head-on collision (**Fig. 2C**). The LEF may translocate past the RNAP at rate *k*_bypass_. However, at any timestep for which LEF remains in the site adjacent to the RNAP and the RNAP translocates toward the LEF, the LEF is pushed by one site along the lattice in the direction of RNAP translocation. If one or more LEF components are on the lattice immediately behind the pushed LEF component, those LEFs are also pushed in the direction of RNAP translocation. In the case where a LEF component is at a lattice site adjacent to the RNAP and the RNAP is translocating *away* from the LEF (*e.g*., they are both moving toward the TTS; **Fig. 2C**), the LEF component may only translocate if it bypasses the RNAP (at rate *k*_bypass_) or if the RNAP vacates (by translocation or unbinding) the lattice site. We focus on results for simulations with characteristic bypassing time *t*_bypass_=1/*k*_bypass_=100 s, *i.e*., we present *k*_bypass_=0.01 (active) and *k*_bypass_=0.0002 (passive), but results for other bypassing rates are shown in **Figs. S4** and **S5**.

## Supplemental Methods

### Generation of HeLa cell lines

All cell lines used in this study were generated by homology-directed repair using CRISPR Cas9 (D10A) paired nickase (107). Based on the cell line SCC1-GFP (11), we introduced a Halo-AID tag to the N-terminus of WAPL, generating Halo-AID-WAPL/SCC1-GFP. Subsequently, Tir1 expression was introduced by transducing a homozygous cell clone with lentiviruses using pRRL containing the constitutive promoter from spleen focus forming virus (SFFV) followed by Oryza sativa Tir1-3xMyc-T2A-Puro (11). Based on this cell line, HA-FKBP12^F36V^-NIPBL/Halo-AID-WAPL/SCC1-GFP and MAU2-FKBP12^F36V^-HA/Halo-AID-WAPL/SCC1-GFP were created by introducing a HA-FKBP12^F36V^ tag to the N terminus of NIPBL or a FKBP12^F36V^-HA tag to the C-terminus of MAU2 individually. As a control cell line, HA-FKBP12 F36V-WAPL was generated using the same gRNAs as described above but with a different repair template. The following gRNAs were used for WAPL: CACCGCTAAGGGTAGTCCGTTTGT and CACCGTGGGGAGAGACCACATTTA; NIPBL: CACCGTCCCCGCAAGAGTAGTAAT and CACCGGTCTCACAGACCGTAAGTT; MAU2: CACCGCTTGGAACTGCACGGGGGG and CACCGCTCCTGTGAGGCCTTGATG. Clones were selected after verification of homozygous integration by PCR of genomic DNA (primers used for WAPL: TGATTTTTCATTCCTTAGGCCCTTG and TACAAGTTGATACTGGCCCCAA; NIPBL: GCAGTGCTTGTCGAGGTTGAT and GCTCAGCCTCAATAGGTACCAACA; MAU2: ATGTCGGTACAGCTGTGGTC and GTGCCACGCACTCTAAGCTA).

### Antibodies and Reagents

Antibodies against MAU2 (Peters laboratory A974) were reported previously (80) and rabbit anti-NIPBL antibodies (Peters laboratory A870 or 133M) were used in a previous publication (37). The following commercial antibodies were used: HA (Abcam, ab9110 for ChIP; BioLegend, 901501 for immunofluorescence and Western blot), GFP (Abcam, ab13970 for immunofluorescence; Abcam, ab290 for ChIP), RNA polymerase II S2 (Abcam, ab5095), SCC1 (Millipore, 05-908), α-tubulin (Sigma-Aldrich, T5168), histone H3 (Cell Signaling, 9715L), and NIPBL (Bethyl, A301-779A). The following secondary antibodies were used: goat anti-chicken IgG Alexa-Fluor-488, anti-mouse IgG Alexa-Fluor-568 (Molecular Probes) for immunofluorescence and anti-rabbit or mouse Ig, and HRP-linked whole antibody (GE Healthcare) for Western blot. Auxin was purchased from Sigma (I5148) and dTAG-7 was a gift from Georg Winter (CeMM, Austria).

### Whole cell extract

Cells were re-suspended in modified RIPA buffer (50 mM Tris pH 7.5, 150 mM NaCl, 1 mM EDTA, 1% NP-40, 0.5% Na-deoxycholate and 0.1% SDS), which additionally contained pepstatin, leupeptin and chymostatin (10 μg/ml each), and PMSF (1 mM). The protein concentration was determined with Bradford assay (Bio-Rad, #5000006). SDS-PAGE and Western Blot were applied to detect individual protein with specific antibodies.

### Chromatin fractionation

Cells were re-suspended with extraction buffer (25 mM Tris pH 7.5, 100 mM NaCl, 5mM MgCl2, 0.2% NP-40, 10% glycerol) supplemented with an EDTA-free protease inhibitor tablet (Roche, 05056489001) and PMSF (1 mM). The chromatin pellet was obtained by centrifugation and re-suspended with extraction buffer containing Benzonase (Merck, 70664) on ice for 30 min, and the protein concentration was determined with Bradford assay.

### Immunofluorescence microscopy

Cells were fixed with 4% formaldehyde for 20 min at room temperature and permeabilized with 0.2% Triton X-100 in PBS for 10 min. After blocking with 3% BSA in PBS-T 0.01% for 30 min, the cells were incubated with the primary antibodies for 1 h and subsequently incubated with the secondary antibodies for 1 h at room temperature. After counterstaining the DNA by 10 min incubation with DAPI, the coverslips were mounted with ProLong Gold (Thermo Fisher) before imaging.

### RNA interference

For RNAi experiments, the cells were transfected as described previously (11). Briefly, the cells were transfected by incubating 100 nM duplex siRNA with RNAi-MAX transfection reagent in antibiotic-free growth medium. After 48 h of RNAi treatment, cells were harvested for experiments. The following target sequences of siRNAs (Ambion) were used: SCC1 (5’-GGUGAAAAUGGCAUUACGGtt −3’), NIPBL (5’-GCAUCGGUAUCAAGUCCCAtt-3’).

### ChIP-seq peak calling and calculation of peak overlaps

Illumina short read sequencing data were aligned against a fusion genome template consisting of merged hg19 and mm9 assemblies. Alignments were filtered for reads which mapped uniquely to the human genome allowing up to two mismatches. The uniquely mappable mouse genome fraction of the spike-in was used for estimation of material loss and calibration by random subsampling of read counts after deduplication.

Peak callings with MACS software versions 1.4 and 2 were applied on full, merged, filtered, and deduplicated replicate data sets of HA-tagged samples with corresponding depleted conditions (+dTAG) as input controls (108). No model-building was performed and output for visualization as bedGraph was normalized by signal strength per million reads for further comparability.

Reference peak set was chosen by searching for these positions in both NIPBL and MAU2 data which could be detected by peak calling via MACS 1.4 using a p-value threshold of 1e-10 in NIPBL data as well as with a p-value threshold of 1e-5 in MAU2 data, and which were confirmed also by MACS2 with default stringencies in both sets. Further, the reference was extended to also include highly scoring NIPBL sites that did not have corresponding signals passing the threshold in the MAU2 fraction (which generally shows weaker signals).

All genomic overlaps as well as area-proportional threefold Venn diagrams have been calculated using multovl version 1.3 (109) and were drawn with eulerAPE (110). Since occasionally more than one site from one dataset overlaps with a single site in a second dataset, resulting coordinates of such an overlap contribute to one single entry - a so-called union. Consequently, the overall site counts drop slightly if displayed in union overlaps.

Heatmap plots have been made using deeptools (111). bedGraph output from peak callings was converted to bigwig input for processing as heatmaps.

RNA polymerase II ChIP-Seq in mouse cells was performed without calibration through spike-in, and short-read sequencing data were mapped against the mm9 template using bowtie2. Only uniquely mappable de-duplicated reads were used further, allowing up to two mismatches.

### Scc1 and CTCF ChIP-seq

ChIP-seq data for Scc1 and CTCF ((6); GSE76303) were mapped using the mm9 assembly and normalized by input (logfe) using a workflow based on the ENCODE pipeline (https://github.com/ENCODE-DCC/chip-seq-pipeline). CTCF sites were selected by overlapping ChIP-seq peaks (6) with CTCF motifs, which were obtained by scanning the position weight matrix (PWM) of the M1 CTCF motif (112) using FIMO (113). Cohesin islands were identified as previously (6).

### Simulations with stationary polymerases

In simulations with stationary RNAP barriers, permeable barriers to extrusion were randomly placed in each gene at the start of the simulation. The probability of placing a barrier at a given polymer site was given by the probability of finding RNAP at that site in steady-state moving barrier simulations. The stationary RNAP simulations therefore modeled a scenario in which all of the RNAP transcribing a gene in steady state suddenly stalled. For the parameters studied, each simulated gene has, at most, a few RNAP. Therefore, to accurately sample the population of genes with stalled RNAP, we performed and averaged approximately 200 simulations.

In these simulations, probability of placing stationary RNAP at the TSS was *p*_TSS_=*k*_load_/(*k*_load_+*k*_unpause_), probability of placing RNAP at a particular gene body site was *p*_body_=*k*_load_*k*_unpause_/(*vp*(*k*_load_+*k*_unpause_)), and probability of placing RNAP at a site beyond the TTS was *p*_tail_(Δ*s*) = *p*_tail,0_ exp{-Δs k_unbind_/v_p_}, where *p*_tail,0_=(*k*_stall_+*k*_unbind_)*k*_load_*k*_unpause_/(*k*_unbind_*v_p_*(*k*_load_+*k*_unpause_)) and Δs is the genomic distance from the polymer site to the 3’ end of the gene.

### Polymer molecular dynamics simulations

3D molecular dynamics simulations with Langevin dynamics were performed with openmm-polymer, as described previously (9, 62, 104), with timestep *dt*=80. *L*=10000 monomeric subunits were arranged into a linear polymer through pairwise harmonic bond interactions. Monomeric subunits repel each other through weak excluded-volume-like interactions.

Each monomeric subunit corresponds to a 1 kb genomic locus simulated in the loop extrusion model. The positions of LEFs indicate additional monomers that are bridged by harmonic bonds. For each configuration of LEFs, the polymer was evolved for 200 timesteps, after which the bonds due to LEFs are updated according to the next step in the extrusion dynamics computation.

Simulations were run for at least 40 LEF residence times, *τ*, before data collection, after which simulations were run for at least 90*τ* At least 4 simulations were performed per parameter set, with 3000 configurations from each simulation included in the analysis. To generate contact maps, we used a cut-off radius *r_c_*=2 monomers, which may be taken to correspond to *r_c_*~50-100 nm (since 1 monomer is 1 kb or a few nucleosomes).

## Data availability statement

Hi-C, Pol II ChIP-seq, and NIPBL ChIP-seq data have been deposited to the Gene Expression Omnibus (GEO), accession number GSE196621. GRO-seq and Scc1 and CTCF ChIP-seq data were previously reported in (6), and they are available on GEO, accession number GSE76303.

## Software availability statement

Codes used for analysis of ChIP-seq and Hi-C data and moving barrier model simulations are freely and publicly available at https://github.com/mirnylab/moving-barriers-paper. Other relevant analysis and simulation codes were previously published and are available as described in the **Materials and Methods** and the **Supplemental Methods**.

## Supplemental Figure Legends

**Supplemental Figure 1.**
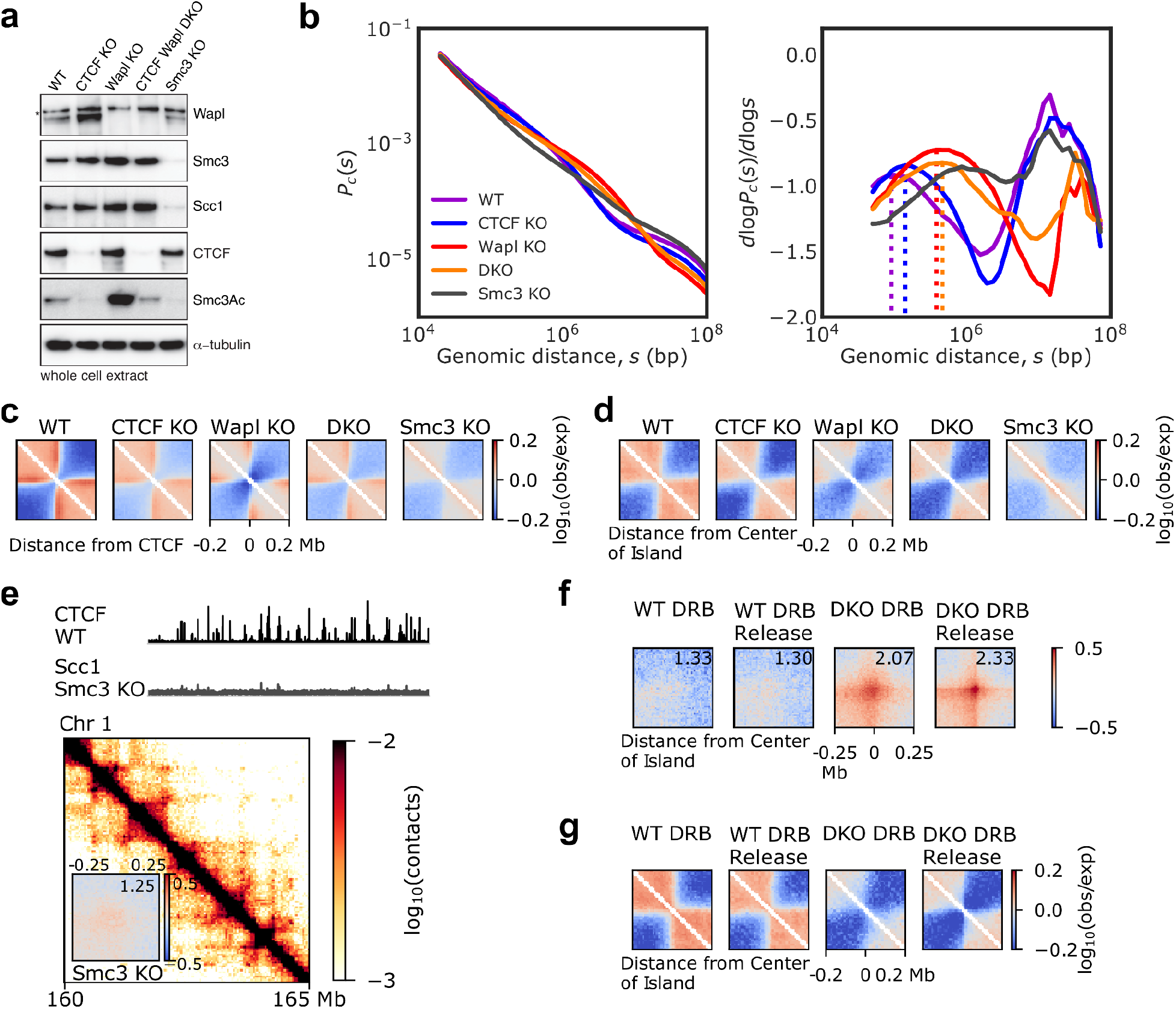
Hi-C in WT, CTCF KO, Wapl KO, DKO, and Smc3 KO cells shows genomic contacts depend on extrusion dynamics. **(A)** Western blots for Wapl, Smc3, Scc1, CTCF, Smc3Ac, and loading control *α*-tubulin in wildtype (control) and different mutants. **(B)** Contact probability, *P_c_*(*s*), and log-derivative, dlog *P_c_*(*s*) / dlog *s*, for WT and mutants, as a function of genomic distance, *s* (left and right, respectively). Dotted lines in log-derivative plot indicate inferred mean loop sizes (64). **(C)** Observed-over-expected maps piled up on the top 10% scoring CTCF sites (*n*=4524), showing insulation around CTCF sites. **(D)** Observed-over-expected maps piled up on previously identified cohesin islands (6). **(E)** Scc1 ChIP-seq and Hi-C contact map of Smc3 KO from the same representative 5 Mb region as in **Fig. 1A**. WT CTCF ChIP-seq track is shown for reference. *Inset*, Observed-over-expected pile ups on island-island dots for Smc3 KO, with dot strength indicated. **(F)** Observed-over-expected pile ups on island-island dots for WT and DKO cells treated with DRB for 3 h and after DRB release by washing 3x with PBS and incubating cells without DRB for 24 h. **(G)** Observed-over-expected pile ups centered on cohesin islands for WT and DKO DRB and DRB release.

**Supplemental Figure 2.**
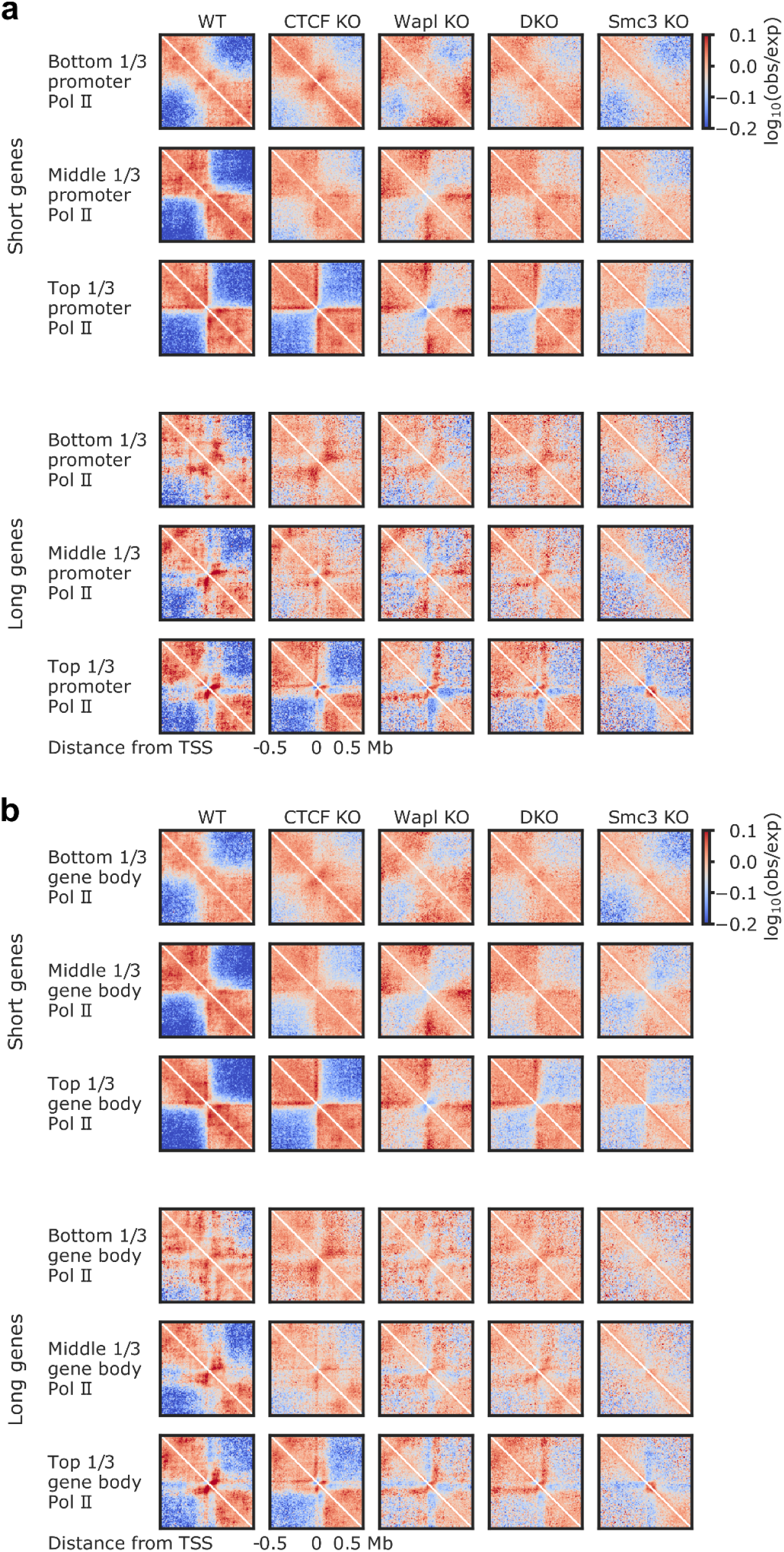
Stratification of genes by RNA polymerase II ChIP-seq signal. **(A)** Average oriented observed-over-expected contact maps for short (10-30 kb; top three rows) and long (80-120 kb; bottom three rows) genes separated into three groups for high, medium, and low RNAP coverage near the TSS. Promoter RNAP was determined by summing RNAP ChIP-seq signal within 2 kb of the TSS. **(B)** Contact maps for short and long genes separated into three groups for high, medium, and low RNAP the entire gene body (taken to be 2 kb upstream of the promoter to 30 kb downstream of the 3’ end of the gene), scaled by gene length.

**Supplemental Figure 3.**
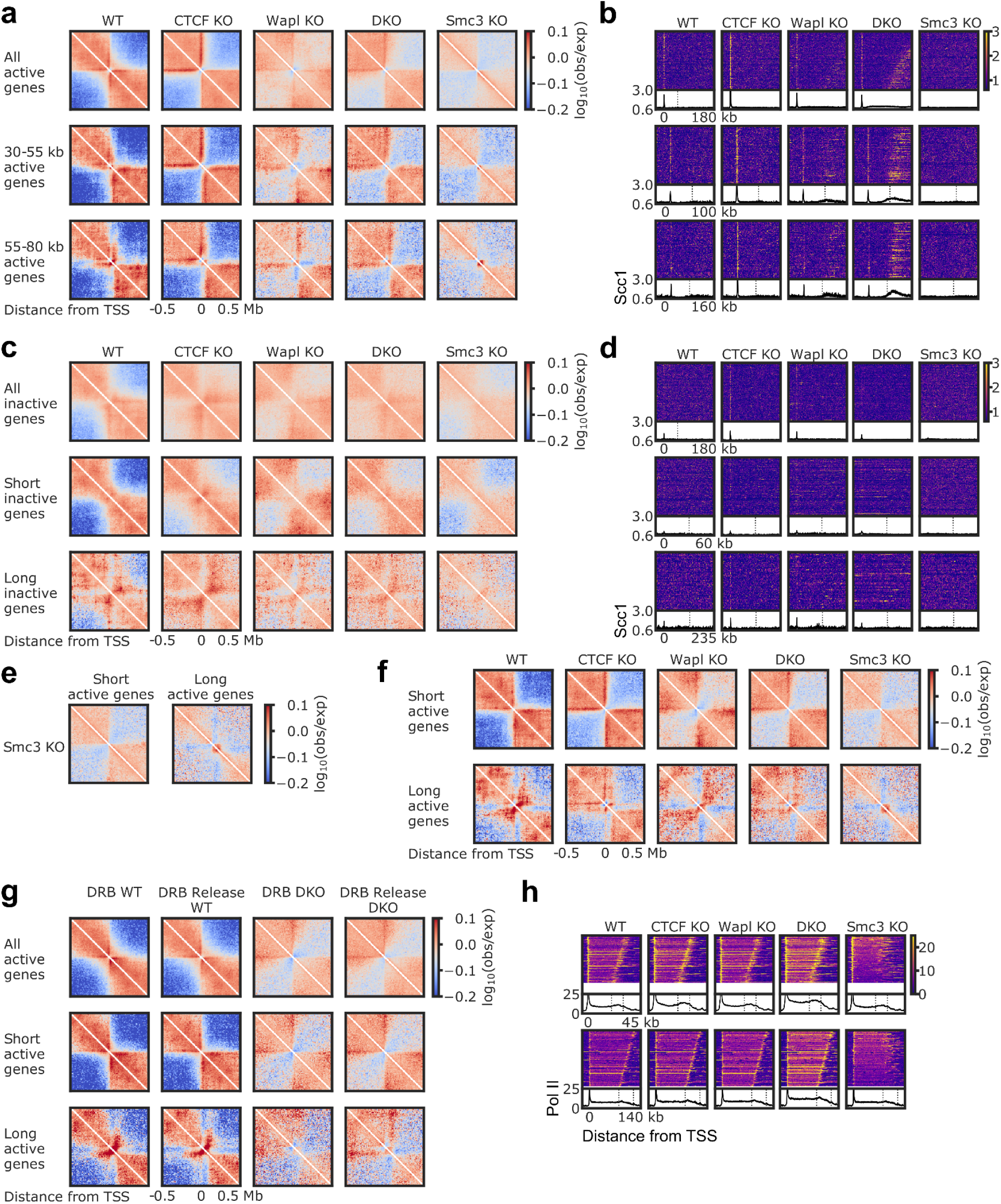
Cohesin dynamics and transcription govern genomic contact patterns near genes. **(A)** Hi-C observed-over-expected pile ups centered on TSSs for all active genes (top row) and two medium-length sets genes (30-55 kb and 55-80 kb, middle and bottom, respectively) for WT, CTCF KO, Wapl KO, DKO, and Smc3 KO. **(B)** Scc1 ChIP-seq heatmaps and average tracks for the same sets of genes. Heat maps show individual gene tracks from the corresponding, sorted by decreasing length from top to bottom, for the longest 50% of genes, except for the set of all active genes, which shows heat maps for the longest 300 genes. Dotted lines show median gene length (49 kb) for all genes or longest gene length for the other two sets. **(C)** Hi-C observed-over-expected pile ups centered on TSSs for all inactive genes (TPM<1, top row), short inactive genes (middle), and long inactive genes (bottom). **(D)** Scc1 ChIP-seq heatmaps and average tracks for the same sets of genes, with heat maps showing the longest 300 genes for the set of all genes and the longest 50% of genes for short and long genes. Dotted lines show median gene length (49 kb) for all genes or longest gene length for the other two sets. **(E)** Pile ups on TSSs for short and long active genes for Smc3 KO. **(F)** Pile ups for short and long active genes for genes that are at least 5 kb away from the top 20% of CTCF sites. **(G)** Pile ups on TSSs for all, short, and long active genes (using TPM>3 from WT GRO-seq or, for DKO, DKO GRO-seq) in WT with DRB treatment or DRB treatment and release and in DKO with DRB treatment or DRB treatment and release. DRB treatment and release were as described in **Fig. S1**. **(H)** RNAP II ChIP-seq heat maps and average tracks for short and long active genes. Heat maps show 300 short genes (here, 20-30 kb) and 168 long genes. All non-overlapping genes were included in RNAP II stack ups. Dotted lines indicate shortest and longest gene lengths in the corresponding sets.

**Supplemental Figure 4.**
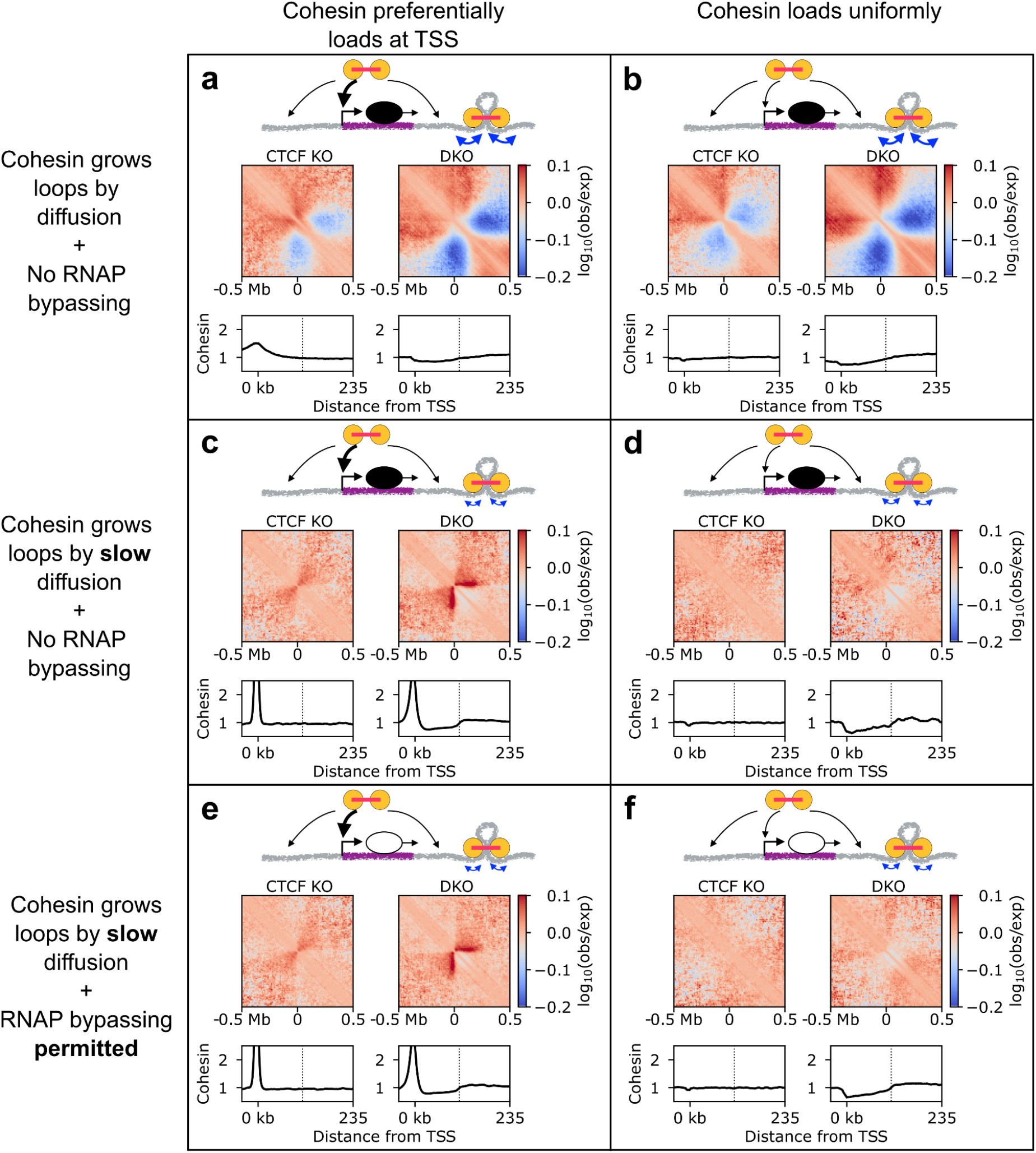
Moving barrier simulations with diffusive cohesins with different diffusion coefficients, RNAP bypassing rates, and loading biases. **(A)** Pile ups centered on TSSs showing observed-over-expected maps for simulations with passive cohesins that grow loops by diffusion and cannot bypass RNAP with loading preferentially at TSSs or **(B)** uniform probability of loading along the genome. **(C)** Simulations with slowly diffusing cohesins (*D*=0.005 kb^2^/s) that cannot bypass RNAP and preferentially load at TSSs or **(D)** load with uniform probability along the genome. **(E)** Slowly diffusing cohesins that *can* bypass RNAPs with cohesin loading preferentially at TSSs or **(F)** cohesin loading uniformly throughout the genome.

**Supplemental Figure 5.**
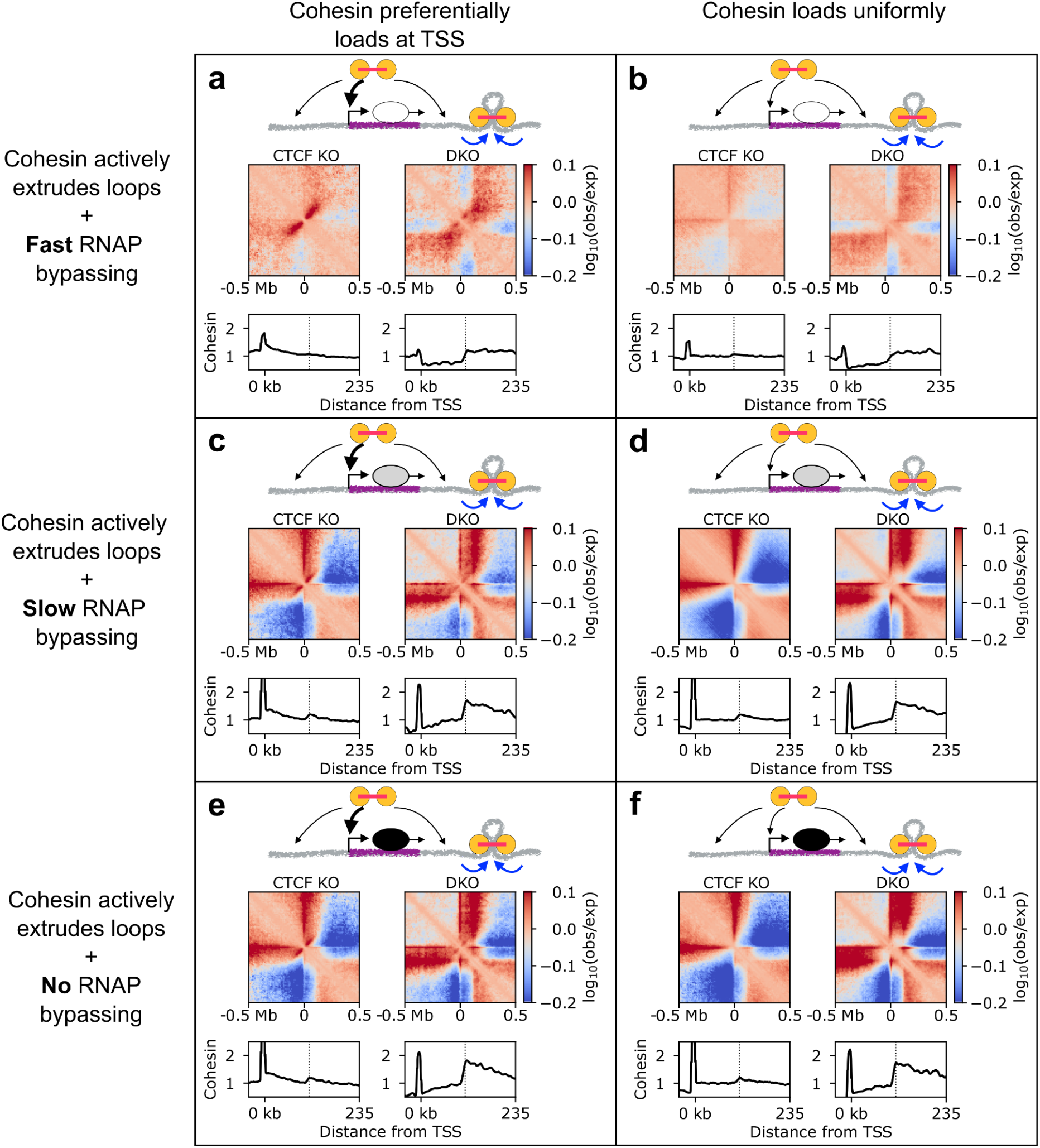
Moving barrier simulations with actively extruding cohesins with different bypassing rates and loading biases. **(A)** Observed-over-expected pile ups and average cohesin accumulation tracks centered on TSSs for active extrusion simulations with fast cohesin-RNAP bypassing (*k*_bypass_=0.04 s^-1^, as opposed to *k*_bypass_=0.01 s^-1^ in the main text) with preferential cohesin loading at TSSs or **(B)** with uniform loading probability across the genome. **(C)** Simulations with active extrusion, slow cohesin-RNAP bypassing (*k*_bypass_=0.001 s^-1^), and preferential cohesin loading at TSSs or **(D)** uniform loading probability throughout the genome. **(E)** Simulations with active extrusion in which cohesin-RNAP bypassing is not permitted (*i.e., k*_bypass_=0 s^-1^) with preferential cohesin loading at TSSs or **(F)** uniform loading.

**Supplemental Figure 6.**
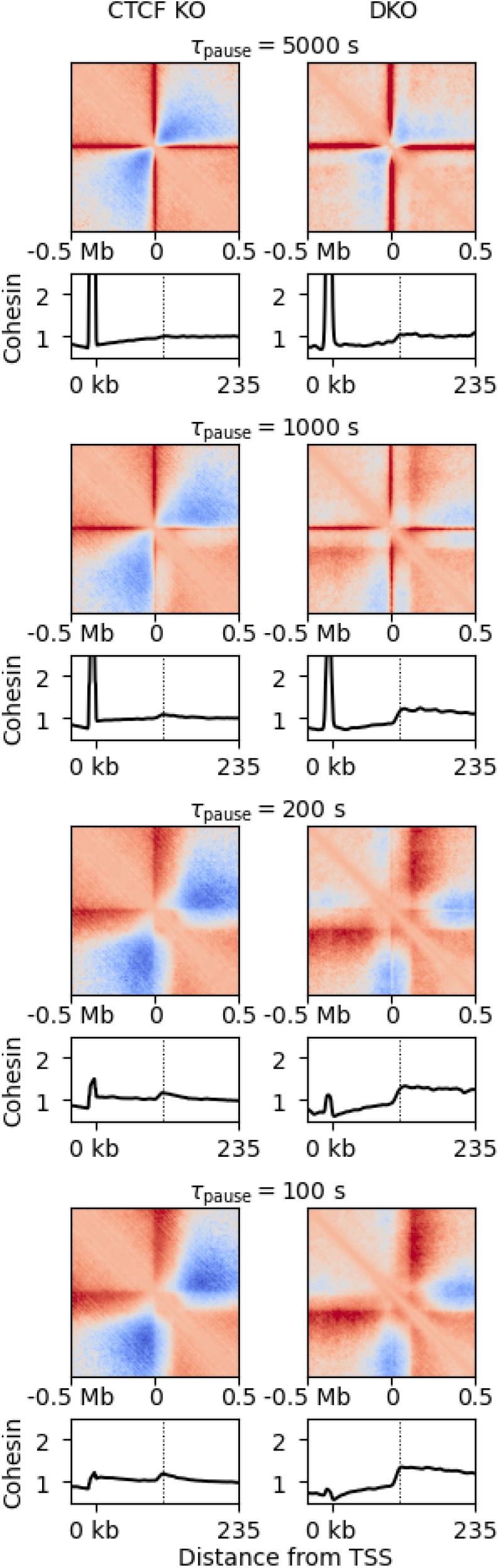
Moving barrier simulations with different RNAP promoter pause times. Observed-over-expected pile ups and average cohesin accumulation tracks centered on TSSs for active extrusion simulations with different RNAP promoter pause times, *τ*_pause_=1/*k*_unpause_. *τ*_pause_ decreases from top row to bottom row. Note *τ*_pause_=500 s in the simulations shown in the main text. Left and right columns, respectively, show CTCF KO and DKO simulations.

**Supplemental Figure 7.**
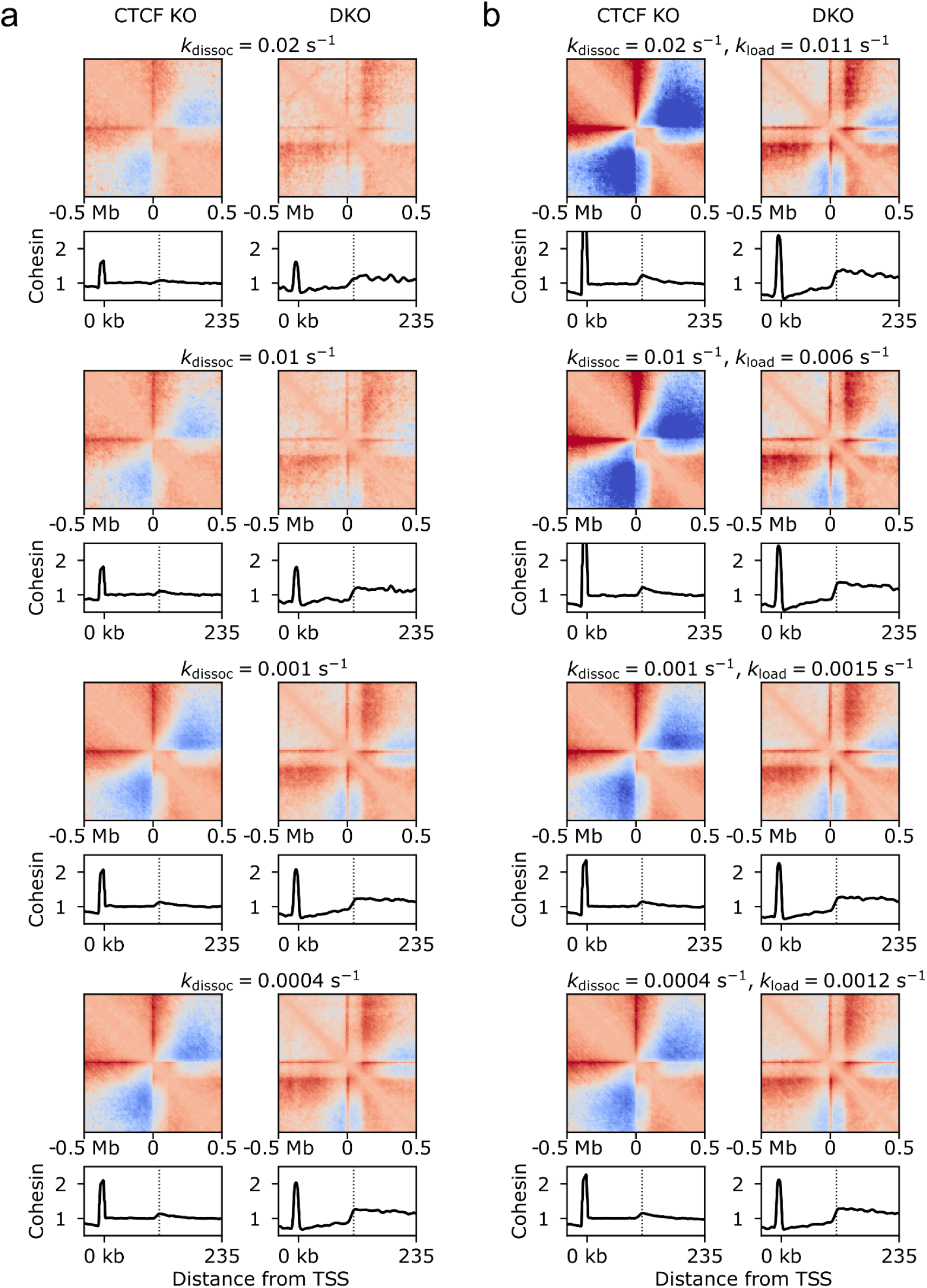
Moving barrier simulations with RNAP that can stochastically dissociate from the promoter. **(A)** Observed-over-expected pile ups and average cohesin accumulation tracks centered on TSSs for active extrusion simulations in which RNAP, while paused at the promoter, can dissociate from chromatin at rate *k*_dissoc_. *k*_dissoc_ decreases from top row to bottom row. Note that RNAP cannot dissociate from chromatin (*k*_dissoc_→∞) in the simulations shown in the main text. Left and right columns, respectively, show CTCF KO and DKO simulations. **(B)** Simulations in which RNAP, while paused at the promoter, can dissociate from chromatin at rate *k*_dissoc_, with RNAP TSS loading rate varied to hold TSS occupancy and level of gene body RNAP constant. *k*_dissoc_ decreases from top row to bottom row. Note that *k*_load_=0.001 s^-1^ in the main text.

**Supplemental Figure 8.**
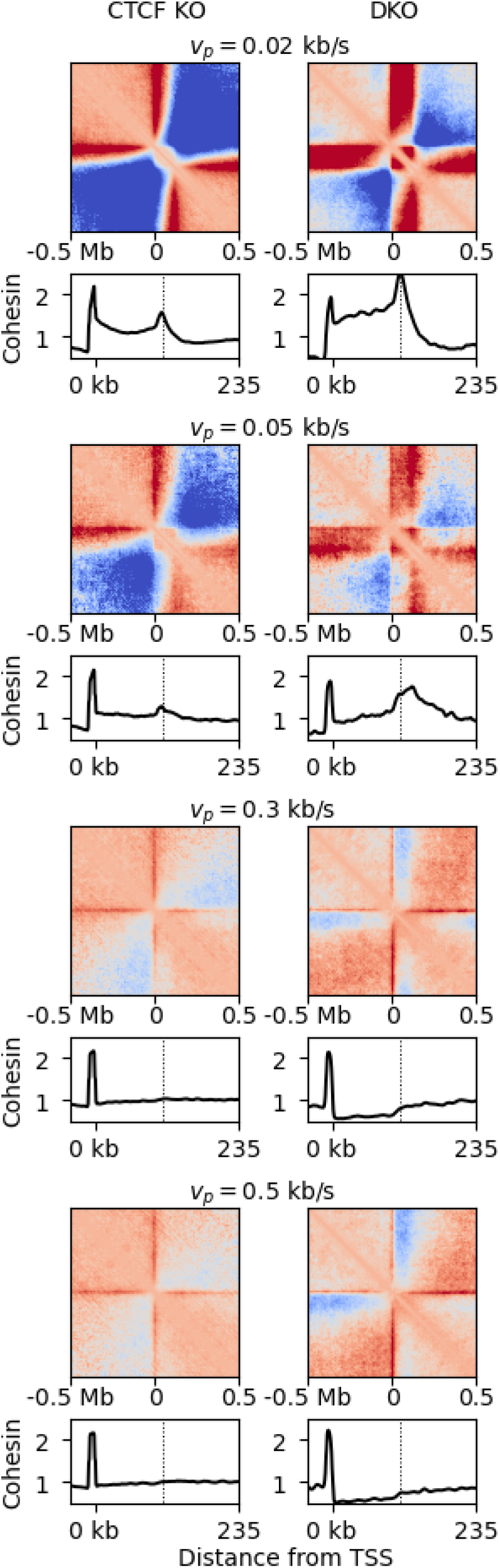
Moving barrier simulations with different RNAP translocation speeds. Observed-over-expected pile ups and average cohesin accumulation tracks centered on TSSs for active extrusion simulations with different RNAP translocation speeds (transcription elongation speeds), *v_p_. v_p_* increases from top row to bottom row. Note *v_p_*=0.1 kb/s for simulations shown in the main text. Left and right columns, respectively, show CTCF KO and DKO simulations. These simulations are performed with a modified configuration of genes on the chromatin polymer (TSSs at *s*=1950, 3950, 5950, 7950 kb) to avoid artifacts in the limit of fast *v_p_*, in which RNAP can overrun the TTS and interfere with the polymer conformation near the neighboring gene.

**Supplemental Figure 9.**
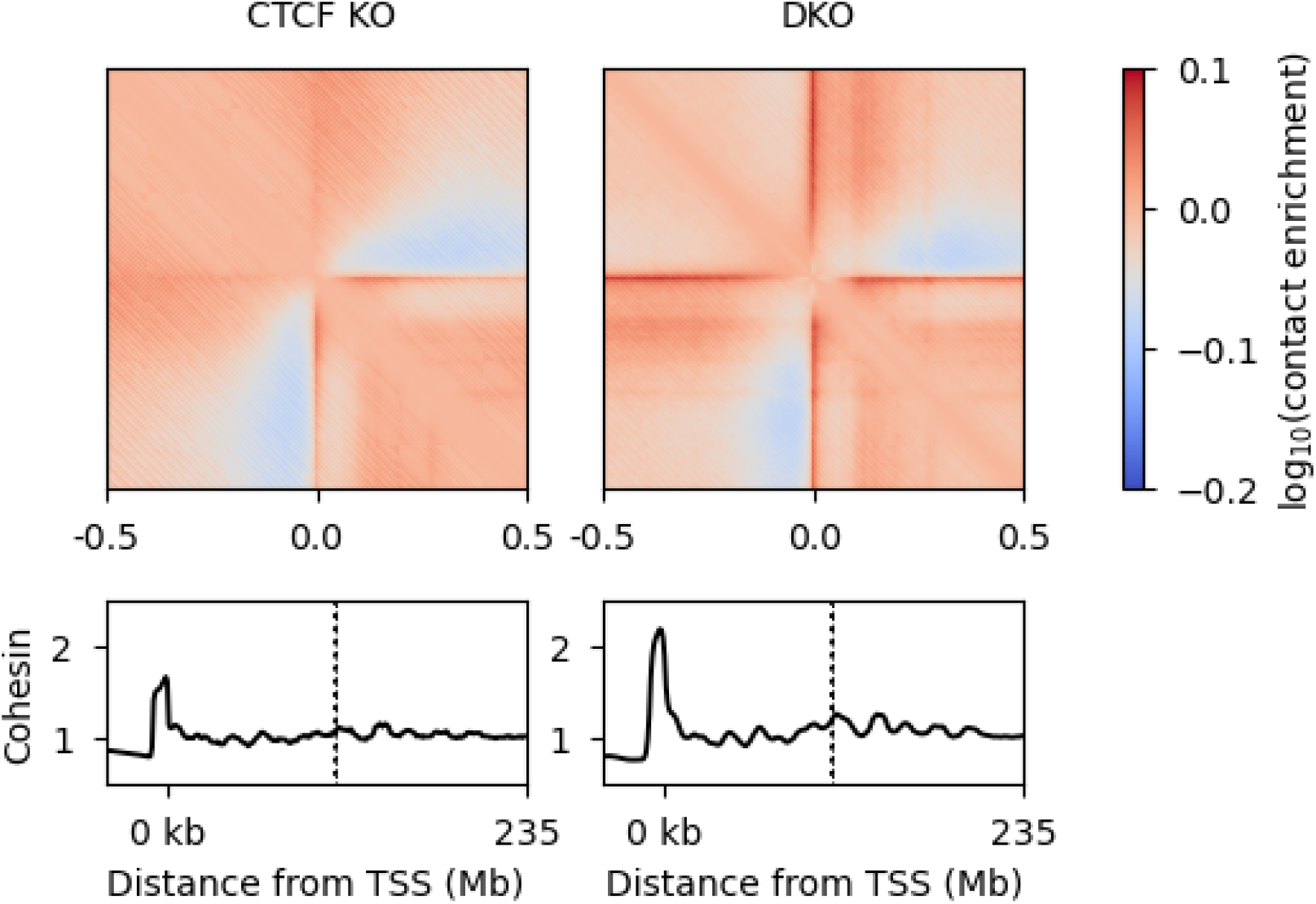
Simulations with RNAPs as stationary barriers distributed randomly throughout genes. Observed-over-expected pile ups and average cohesin accumulation tracks centered on TSSs for active extrusion simulations RNAP distributed as stationary barriers throughout the TSS, gene, and region downstream of the 3’ end of the gene (see **Materials and Methods**). Left and right columns, respectively, show CTCF KO and DKO simulations.

**Supplemental Figure 10.**
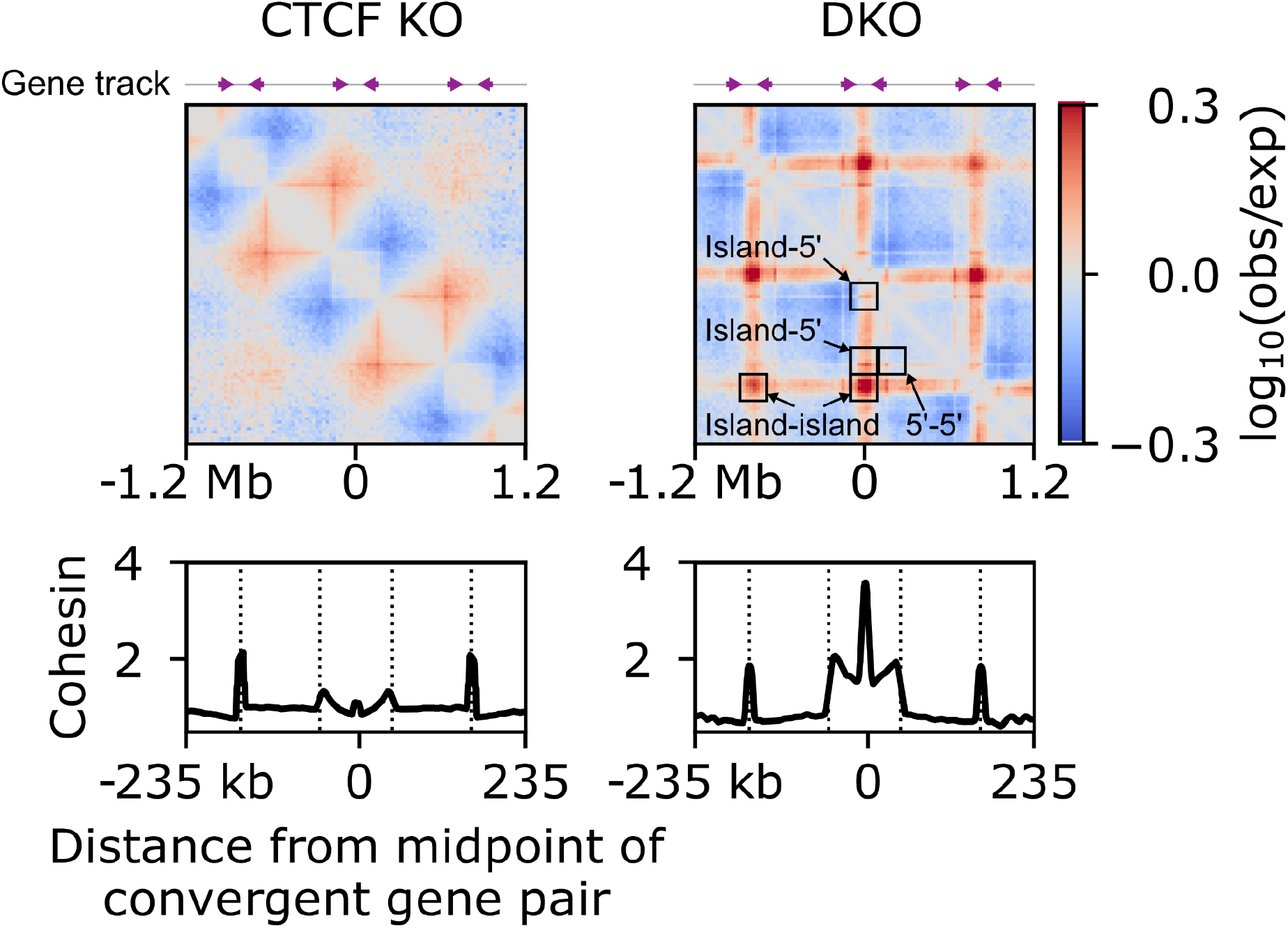
Simulations of convergently oriented genes produce cohesin islands and island-island dots in DKO conditions. *Top*, Pile ups centered on sites of convergent transcription showing observed over expected contacts for CTCF KO and DKO simulations. Gene track (top) indicates positions, lengths, and orientations of genes (purple arrows) on the chromosome polymer fiber (gray). Boxes identify features of interest, such as contacts formed between cohesin islands or contacts between gene ends. *Bottom*, Average cohesin occupancy along the polymer fiber in simulations in a small region around a convergent gene pair. Dotted lines indicate the 5’ and 3’ ends of genes.

**Supplemental Figure 11.**
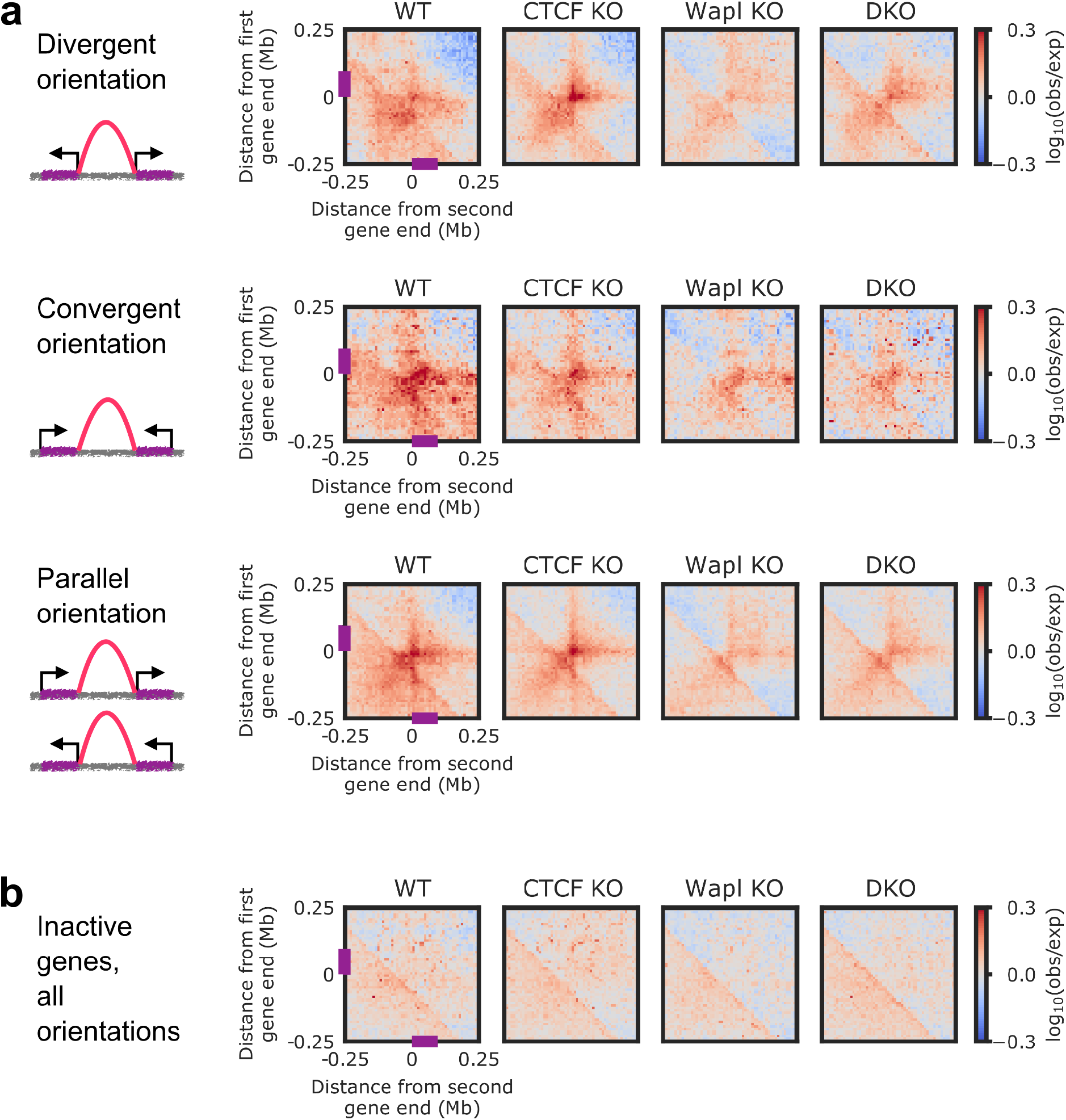
3’ and 5’ ends of active genes act as extrusion barriers and generate genomic contacts. **(A)** Pile ups centered on contacts between nearest ends of pairs of adjacent active genes (TPM>2) within a genomic distance 50 kb< *s* < 350 kb of each other, separated by relative orientation: convergent, divergent, and parallel (top, middle, and bottom, respectively). Schematic drawings illustrate the orientations. Purple bars indicate positions of genes in the pile ups. **(B)** Pile ups centered on contacts of nearest ends of pairs of adjacent inactive genes (TPM<1) within a genomic distance 50 kb< *s* < 350 kb of each other, where at least one gene in each pair is not near a CTCF site (see **Materials and Methods**).

**Supplemental Figure 12.**
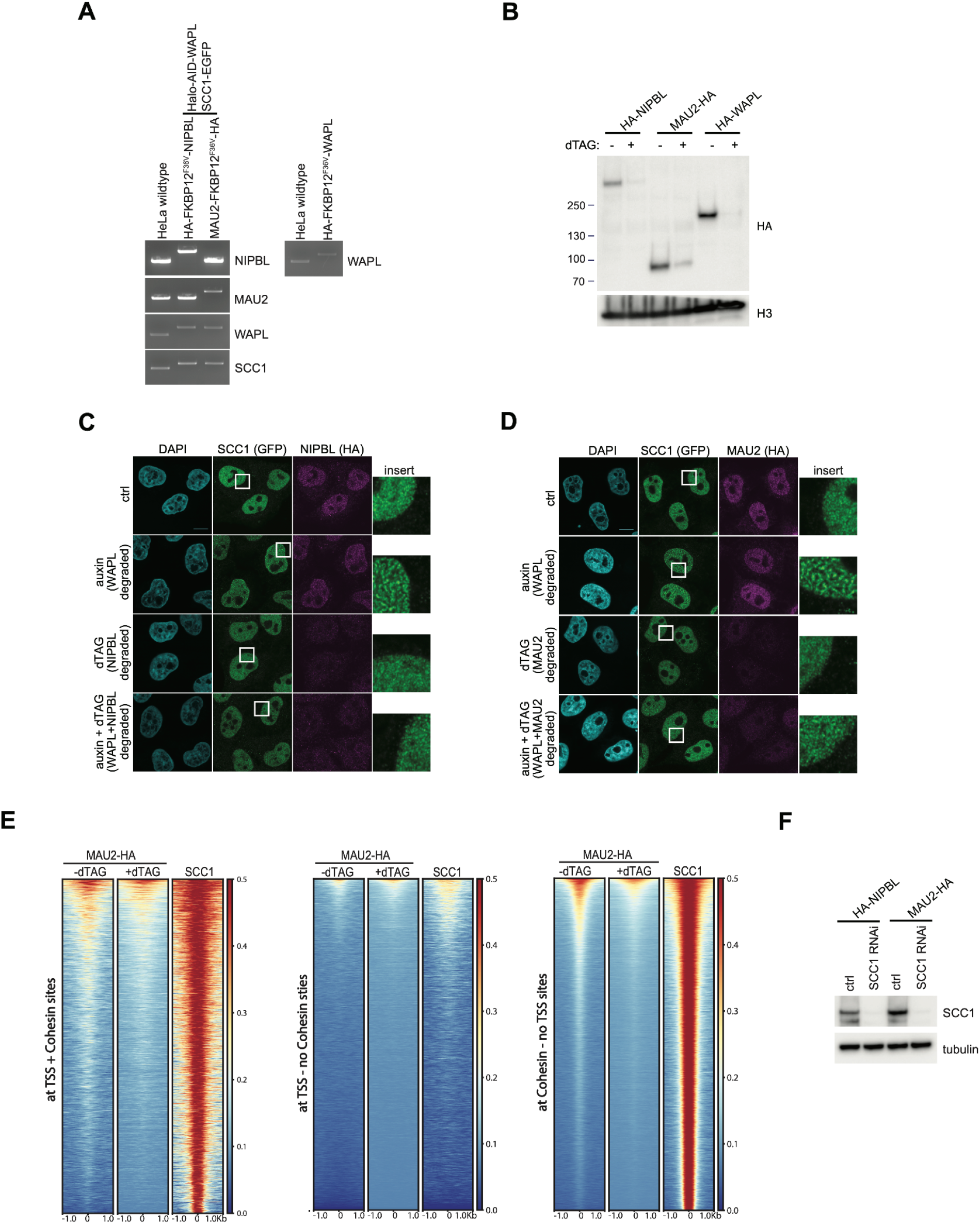
NIPBL and MAU2 colocalize predominantly with cohesin but not TSSs throughout the genome. **(A)** Genotype analysis of HeLa wildtype, homozygous HA-FKBP12^F36V^-NIPBL/Halo-AID-WAPL/SCC1-EGFP (HA-NIPBL), MAU2-FKBP12^F36V^-HA/ Halo-AID-WAPL/SCC1-EGFP (MAU2-HA) and HA-FKBP12^F36V^-WAPL (HA-WAPL) cells. **(B)** Chromatin fractionation of HA-NIPBL, MAU2-HA and HA-WAPL cells were analyzed by immunoblotting with or without dTAG for 24 hours. **(C)** Representative immunofluorescence images of HA-NIPBL cells treated with auxin or/and dTAG for 24 hours. DNA was counterstained with DAPI. *Inset*, magnified images of boxed regions. Scale bar shows 10 μm. **(D)** Representative immunofluorescence images of MAU2-HA cells treated with auxin or/and dTAG for 24 hours. DNA was counterstained with DAPI. *Inset*, magnified images of boxed regions. Scale bar shows 10 μm. **(E)** Heatmaps of MAU2-HA (-/+dTAG) and SCC1 ChIP-Seq signal at TSSs with cohesin, TSSs without cohesin, and cohesin without TSSs. **(F)** Immunoblotting analysis of HA-NIPBL or MAU2-HA cells treated with or without SCC1 siRNA for 48 hours.

**Supplemental Figure 13.**
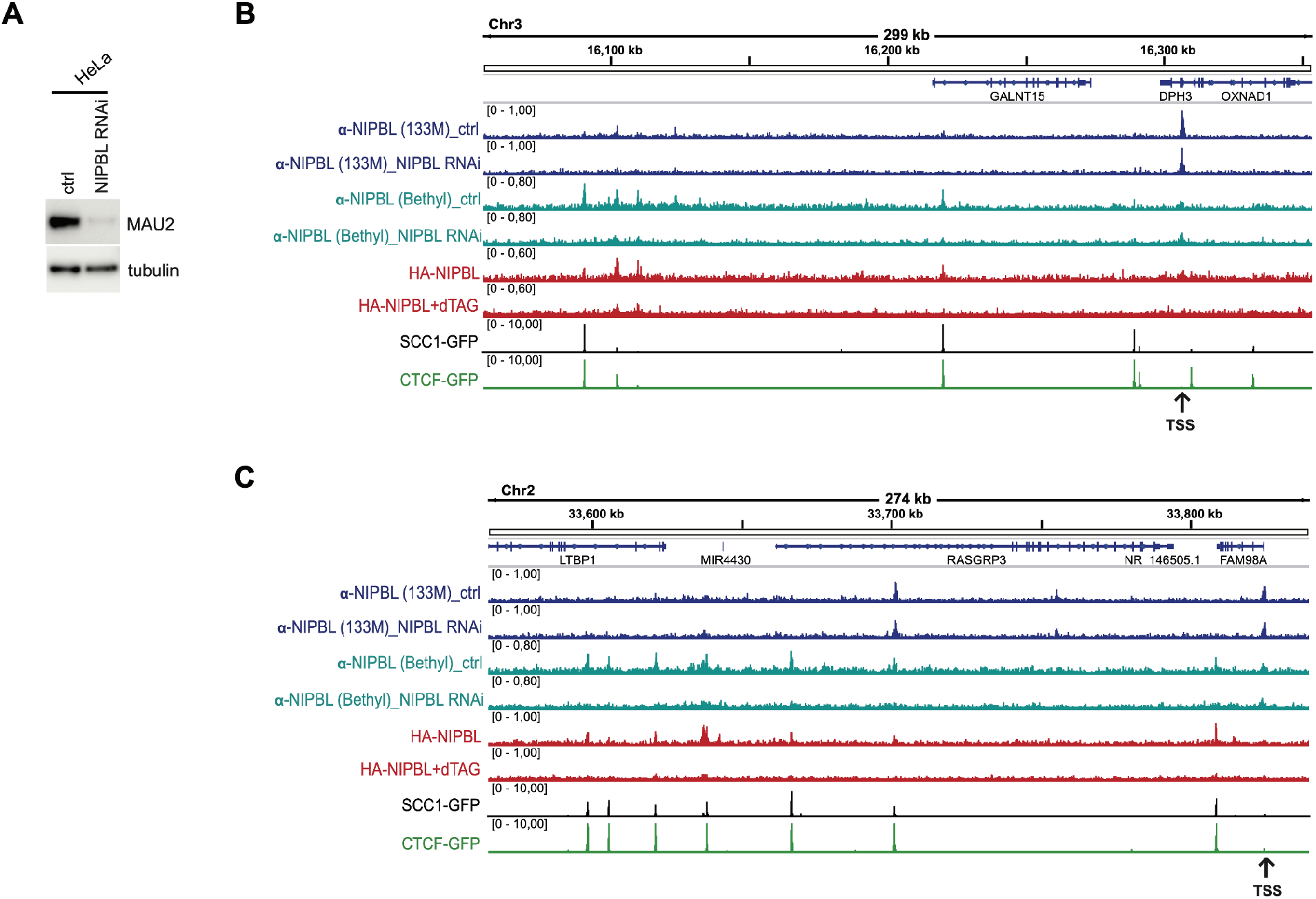
ChIP-seq profiles from three different NIPBL antibodies. **A)** Immunoblotting analysis of HeLa cells treated with or without NIPBL siRNA for 48 hours. **(B)** Enrichment profiles of NIPBL (133M and Bethyl antibody, -/+ NIPBL RNAi), HA-NIPBL (-/+dTAG), SCC1-GFP and CTCF-GFP along an exemplary 299 kb region of chromosome 3, illustrating typical distribution and co-localization of sequencing read pileups. Genes within this region are depicted above. The arrow beneath indicates the transcription start site. **(C)** Enrichment profiles of NIPBL (133M and Bethyl antibody, -/+ NIPBL RNAi), HA-NIPBL (-/+dTAG), SCC1-GFP and CTCF-GFP along an exemplary 274 kb region of chromosome 2, illustrating typical distribution and co-localization of sequencing read pileups. Genes within this region are depicted above. The arrow indicates the transcription start site.

**Supplemental Figure 14.**
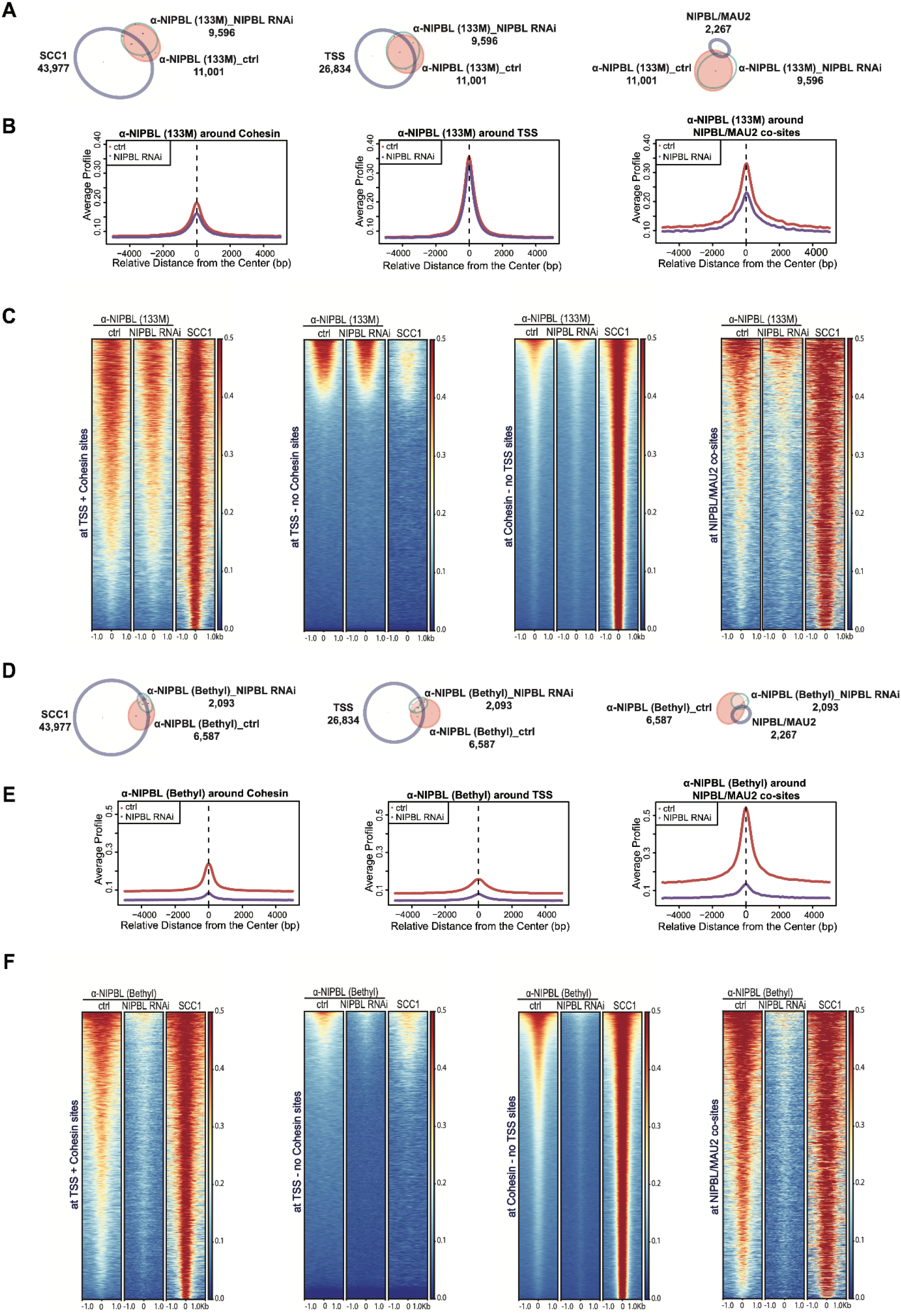
Bioinformatic analysis of ChIP-seq data from three different NIPBL antibodies. **(A)** Area-proportional threefold eulerAPE Venn diagram illustrating overlap between NIPBL (133M, -/+ NIPBL RNAi) and SCC1, TSSs, or NIPBL/MAU2 co-sites (left, middle, and right, respectively). **(B)** Average signal profiles of NIPBL (133M, -/+ NIPBL RNAi) binding around cohesin, TSSs or NIPBL/MAU2 co-sites. **(C)** Heatmaps of NIPBL (133M, -/+ NIPBL RNAi) and SCC1 at TSSs with cohesin, TSSs without cohesin, cohesin sites without TSSs, or NIPBL/MAU2 co-sites. **(D)** Area-proportional threefold eulerAPE Venn diagram illustrating overlap between NIPBL (Bethyl antibody, -/+ NIPBL RNAi) and SCC1, TSSs, or NIPBL/MAU2 co-sites (left, middle and right, respectively). **(E)** Average signal profiles of NIPBL (Bethyl antibody, -/+ NIPBL RNAi) binding around cohesin, TSSs or NIPBL/MAU2 co-sites. **(F)** Heatmaps of NIPBL (Bethyl antibody, -/+ NIPBL RNAi) and SCC1 at TSSs with cohesin, TSSs without cohesin, cohesin sites without TSSs, or NIPBL/MAU2 co-sites.

**Supplemental Figure 15.**
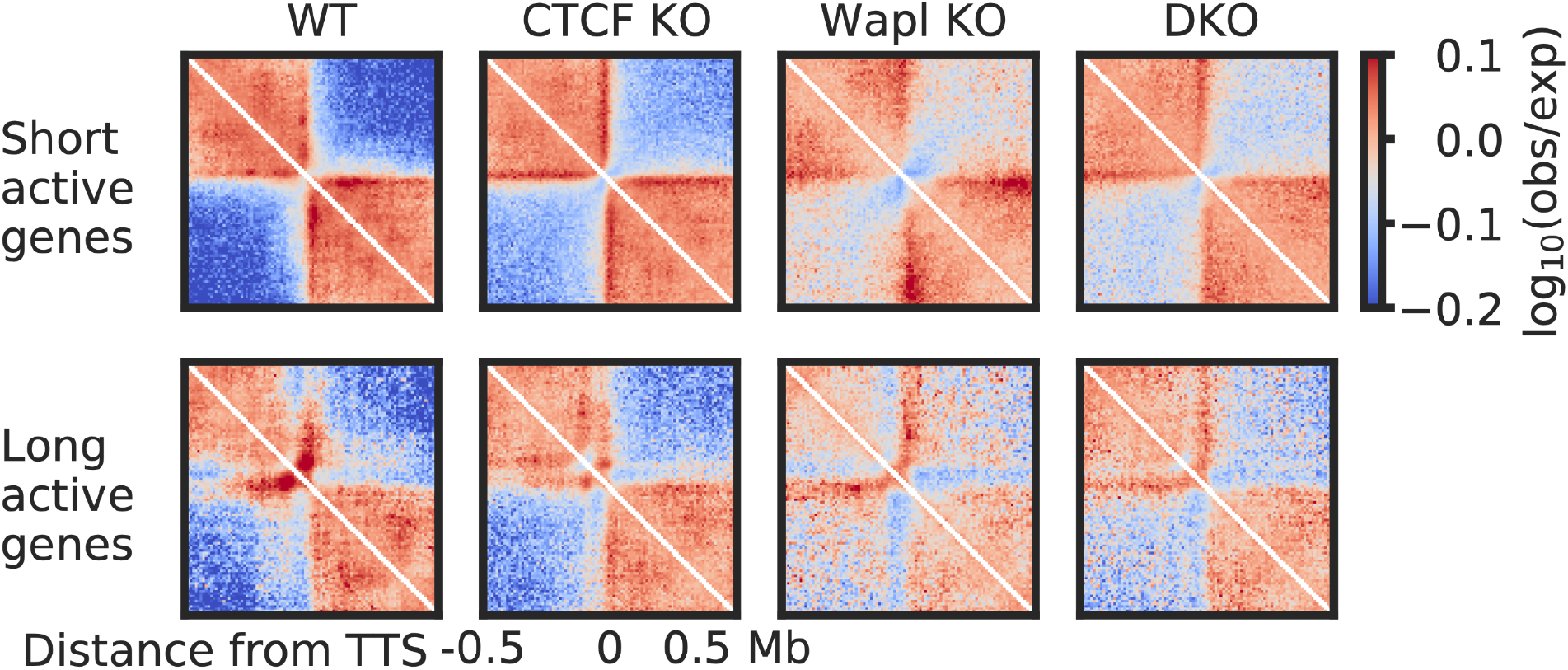
Transcription termination sites of active genes generate insulation. Observed over expected Hi-C contact maps for short and long active genes in WT, CTCF KO, Wapl KO, and DKO, piled up and centered on their transcription termination sites (TTSs).

## Acknowledgements

We thank Nezar Abdennur, Sameer Abraham, Anton Goloborodko, and Maxim Imakaev for instructive discussions and sharing code. We also thank Job Dekker, Anne-Laure Valton, and Sergey Venev for helpful conversations. We are grateful to the Sequencing Team at IMP Vienna for their assistance in the sequencing experiments. HBB was partially supported by the National Institutes of Health (NIH) grant R00GM130896. LAM acknowledges funding from R01GM114190 of the NIH, the NIH Common Fund 4D Nucleome Program (UM1HG011536), the Human Frontier Science Program (grant RGP0057/2018), and Chaire Internationales d’excellence Blaise Pascal. JMP acknowledges funding from Boehringer Ingelheim, the Austrian Research Promotion Agency (Headquarter grant FFG-852936), the European Research Council (ERC) under the European Union’s Horizon 2020 research and innovation programme (grant agreements No 693949 and No 101020558), the Human Frontier Science Program (grant RGP0057/2018), and the Vienna Science and Technology Fund (grant LS19-029). For the purpose of Open Access the authors have applied a CC-BY public copyright license to any Author Accepted Manuscript version arising from this submission. JMP is also an adjunct professor at the Medical University of Vienna.

## Author Contributions

EJB, AAB, HBB, and LAM conceived and designed the simulation model. EJB and AAB performed and analyzed the simulations. LAM supervised the simulation research. WT, GW, GAB, and JMP conceived and designed the experiments. WT and GW performed the experiments. EJB, WT, AAB, RRS, and GW analyzed the experimental data. JMP supervised the experimental research. LAM and JMP supervised analysis of the experimental data. EJB drafted the manuscript with input from LAM and JMP. All authors discussed the results and commented on the manuscript.

